# A class IV adenylate cyclase CyaB is required for capsule polysaccharide production and biofilm formation in *Vibrio parahaemolyticus*

**DOI:** 10.1101/2022.06.06.495059

**Authors:** A. Regmi, J.G. Tague, K. Boas Lichty, E.F. Boyd

**Affiliations:** Department of Biological Sciences, University of Delaware, Newark, DE 19716

**Author notes:** Corresponding author, E. Fidelma Boyd, Department of Biological Sciences, 341 Wolf Hall, University of Delaware Newark, DE 19716.

## Abstract

CRP (cyclic AMP receptor protein), encoded by *crp*, is a global regulator that is activated by cAMP (cyclic AMP), a second messenger synthesized by a class I adenylate cyclase (AC-I) encoded by *cyaA* in *Escherichia coli*. cAMP-CRP is required for growth on non-preferred carbon sources and is a global regulator. We constructed in-frame non-polar deletions of the *crp* and *cyaA* homologs in *Vibrio parahaemolyticus* and found that the Δ*crp* mutant did not grow in minimal media supplemented with non-preferred carbon sources, but the Δ*cyaA* mutant grew similar to wild type. Bioinformatics analysis of the *V. parahaemolyticus* genome identified a 181 amino acid protein annotated as a class IV adenylate cyclase (AC-IV) named CyaB, a member of the CYTH protein superfamily. AC-IV phylogeny showed CyaB was present in Gamma- and Alpha- Proteobacteria as well as Planctomycete and Archaea. Only the bacterial CyaB proteins contained an N-terminal motif HFxxxExExK indicative of adenylyl cyclase activity. Both *V. parahaemolyticus cyaA* and *cyaB* genes functionally complemented an *E. coli* Δ*cyaA* mutant. The Δ*crp* and Δ*cyaB/*Δ*cyaA* mutants showed defects in growth on non-preferred carbon sources, and in swimming and swarming motility, indicating cAMP-CRP is an activator. The Δ*cyaA* and Δ*cyaB* single mutants had no defects in these phenotypes indicating AC-IV complements AC-I. Capsule polysaccharide and biofilm production assays showed significant defects in Δ*crp*, Δ*cyaB/*Δ*cyaA,* and the Δ*cyaB* mutant, whereas Δ*cyaA* behaved similar to wild type. This is consistent with a role of cAMP-CRP as an activator of these phenotypes and establishes a cellular role for AC-IV in capsule and biofilm formation, which to date has been unestablished.

**IMPORTANCE:** Here, we characterized the roles of CRP and CyaA in *V. parahaemolyticus,* showing cAMP-CRP was an activator of metabolism, motility, capsule and biofilm formation. These results are in contrast to cAMP-CRP in *V. cholerae,* which represses capsule and biofilm formation. Previously, only an AC-I CyaA had been identified in *Vibrio* species. Our data showed that an AC-IV CyaB homolog is present in *V. parahaemolyticus* and was required for optimal growth. The data demonstrated that CyaB was essential for capsule production and biofilm formation uncovering a physiological role of AC-IV in bacteria. The data showed that the *cyaB* gene was widespread among *Vibrionaceae* species and several other Gamma-Proteobacteria, but in general, its phylogenetic distribution was limited. Our phylogenetic analysis also demonstrated that in some species the *cyaB* gene was acquired by horizontal gene transfer.

## INTRODUCTION

Carbon catabolite repression (CCR) is a global regulatory mechanism that represses functions required for secondary carbon metabolism when a preferred carbon source is present (1, 2). In *Escherichia coli,* CCR is mediated by several mechanisms such as the regulation of the biosynthesis of catabolic enzymes, the inhibition of carbon transport, and metabolic signaling (3, 4). Some of the major players involved in CCR are the global regulator CRP (**c**AMP **R**eceptor **P**rotein), adenylate cyclase (AC), the signaling molecule cAMP (3’,5’-cyclic AMP), and components of phosphoenolpyruvate:sugar phosphotransferase system (PTS) transporters (1, 3–7). CRP functions as a homodimer, activated by the second messenger cAMP and in *E. coli* cAMP-CRP is a global transcriptional activator of non-preferred carbon genes in the absence of glucose (8–10). In *E. coli*, CRP has an N-terminal cAMP binding domain (CBD; residues 1–136) and a C- terminal helix turn helix DNA binding domain (DBD; residues 139–209) (11, 12). When bound by cAMP, CRP undergoes a conformation change that allows the protein to interact with target promoter regions at consensus sequences (13, 14).

When glucose is present, it is imported by the PTS transporter, resulting in dephosphorylation of the EIIA component of PTS. EIIA^glc^ inhibits non-preferred carbon transporters, a mechanism known as inducer exclusion, independent of cAMP signaling (2, 4, 15–17). When glucose is absent, EIIA^glc^ is phosphorylated and P∼EIIA^glc^ activates adenylate cyclase to produce cAMP (4, 18). However, growth on non-PTS carbon sources, such as lactose, and in limited nitrogen and phosphorous conditions also have reduced cAMP levels, indicating the presence of a catabolite modulating factor, which Monod suggested in his last paper (19). Recent research showed that cAMP signals were modulated by metabolic precursors such as α-ketoacids to control the allocation of proteomic resources by a feedback system rather than carbon metabolism (20, 21).

There are six classes (I-VI) of adenylate cyclases (AC) based on primary amino acid sequence, all with different catalytic mechanisms and structures, indicating different evolutionary origins (22, 23). *Escherichia coli* and many other *Proteobacteria* contain a single AC gene *cyaA*, which encodes a class I AC (AC-I). The 810 amino acid CyaA protein has two functionally distinct domains, the N-terminal catalytic cyclase domain and the C-terminal regulatory domain (11, 12, 24). Class II ACs (AC-II) are comprised of bacterial toxins such as the exotoxin from *Bordetella pertussis* (25). The class III ACs (AC-III) are the most diverse and widespread, and encompass adenylate cyclases present in eukaryotes and prokaryotes (26). A structurally diverse AC was identified in *Prevotella ruminicola* D31d and forms the sole representative of AC-V (27). A class VI AC was described in *Rhizobium elti*, a 345 amino acid protein that was shown to have cyclase activity by complementation of an *E. coli cyaA* deletion mutant (28).

Class IV AC (AC-IV) along with mammalian thiamine triphosphatases and inorganic triphosphatases belong to the CYTH-like domain superfamily (29). CYTH proteins are present in all three domains, Bacteria, Archaea and Eukaryota, and perform a variety of functions (29). CYTH proteins contain a triphosphate tunnel metaloenzyme (TTM) fold and act on triphosphorylated substrates with at least one divalent metal cation required for catalysis. A hallmark of CYTH proteins is an N-terminal ExExK motif (30). In bacteria, an AC-IV CyaB was initially described in *Aeromonas hydrophila,* a small 181 amino acid protein with a single domain (31). The foundational work in *A. hydrophila* indicated that *cyaB* was not expressed under standard growth conditions, but *cyaB* did functionally complement an *E. coli cyaA* deletion mutant (31). A CyaB homolog was also identified from *Yersinia pestis*, its crystal structure (2FJT) was determined and showed to be very different from AC-I and AC-III enzymes. AC-IV formed a dimer like AC-III, but with a distinct antiparallel eight-stranded barrel active site and could cyclize both ATP and GTP (32–34). The physiological role of bacterial AC-IV was not established. Archaea contain numerous proteins annotated as AC-IV, but cAMP biosynthesis has not been shown for this domain (35). A recent study examining a CYTH-like protein annotated as an AC-IV from the Crenarchaeote *Sulfolobus acidocaldarius* showed that it had triphosphatase activity, but no adenylate cyclase activity (35). Vogt and colleagues differentiated CYTH-like proteins (with phosphatase activity) from AC-IV (with adenylyl cyclase activity) using sequence similarity network analysis and functional studies (35). They suggested that a true AC-IV has an extended N-terminal motif HFxxxxExExK and the absence of HF indicates a CYTH protein with either thiamine triphosphatase (ThTPase) or inorganic triphosphatase activity (35).

In the present study, the function and activation of CRP in *V. parahaemolyticus* was investigated for the first time. In-frame non-polar deletions of *crp* (VP2793) and *cyaA* (VP2987) in *V. parahaemolyticus* RIMD2201633 were constructed and examined for growth on various carbon sources. The Δ*crp* mutant grew on minimal medium M9 3% NaCl supplemented with glucose (M9G) and was unable to grow on non-PTS carbon sources, but the Δ*cyaA* mutant grew similar to wild type. This suggested the presence of an additional uncharacterized AC in *V. parahaemolyticus*. Bioinformatics analysis identified ORF VP1760 that encoded a putative AC-IV named CyaB. The phylogeny of AC-IV among Bacteria and their closest orthologs in Archaea was examined to determine its distribution and evolutionary history. To examine whether *cyaA* and *cyaB* were both functional, an *E. coli* Δ*cyaA* mutant was functionally complemented with both genes. Growth pattern analysis of the Δ*cyaB,* Δ*cyaA* and a double Δ*cyaB/*Δ*cyaA* mutants on different carbon sources was also examined. Swimming and swarming motility assays, capsule polysaccharide (CPS) production and biofilm formation assays were performed to determine whether cAMP-CRP played a role in these important phenotypes.

## RESULTS

### CRP is an essential regulator in *V. parahaemolyticus*

cAMP-CRP is a global regulator of nutrient transport and metabolism in many bacterial species (13, 36–40). Homologs of CRP and CyaA are present in *V. parahaemolyticus*, ORF VP2793 and VP2987, respectively (**Fig. S1**). The CRP and CyaA proteins of *V. parahaemolyticus* shared 98% and 82% amino acid identity respectively to homologs in *V. cholerae*. The genome context of both genes is conserved across *V. parahaemolyticus*, *V. cholerae,* and *E. coli* (**Fig. S1**). To begin to investigate the physiological role of CRP in *V. parahaemolyticus,* we first constructed an in-frame non-polar deletion of *crp* (VP2793) and *cyaA* (VP2987) in *V. parahaemolyticus* RIMD2210633. The Δ*crp* mutant displayed a small colony morphology on LB supplemented with 3% NaCl (LBS) plates **(Fig. S2A)**.

Growth pattern analysis in LBS broth and M9S D-glucose broth demonstrated that the Δ*crp* strain had a growth defect **(Fig. 1A**-**B**). Next, we examined the Δ*cyaA* mutant, expecting a phenotype similar to the Δ*crp* mutant, as shown in *E. coli* and *V. cholerae* Δ*cyaA* mutants. However, growth of Δ*cyaA* on LBS plates gave identical colony morphology to wild type (**Fig. S2A)** and similarly in LBS and M9 supplemented with 3% NaCl (M9S) and D-glucose broth, Δ*cyaA* presented a similar pattern to wild type (**Fig. 1A-B**). In addition, growth in alternative carbon sources, M9S supplemented with L-arabinose, L- ribose, or D-gluconate, showed the Δ*crp* strain was defective, but the Δ*cyaA* mutant grew similar to wild type (**Fig 2A-C**). These data demonstrate that CRP is required for catabolism of alternative carbon sources similar to *V. cholerae*, but that *V. parahaemolyticus* contained an additional uncharacterized adenylate cyclase that can synthesize cAMP in the absence of *cyaA* to activate CRP. Alternatively, CRP could be acting independently of cAMP, but given its high homology to CRP from *V. cholerae* this is less likely.

**Fig. 1.**
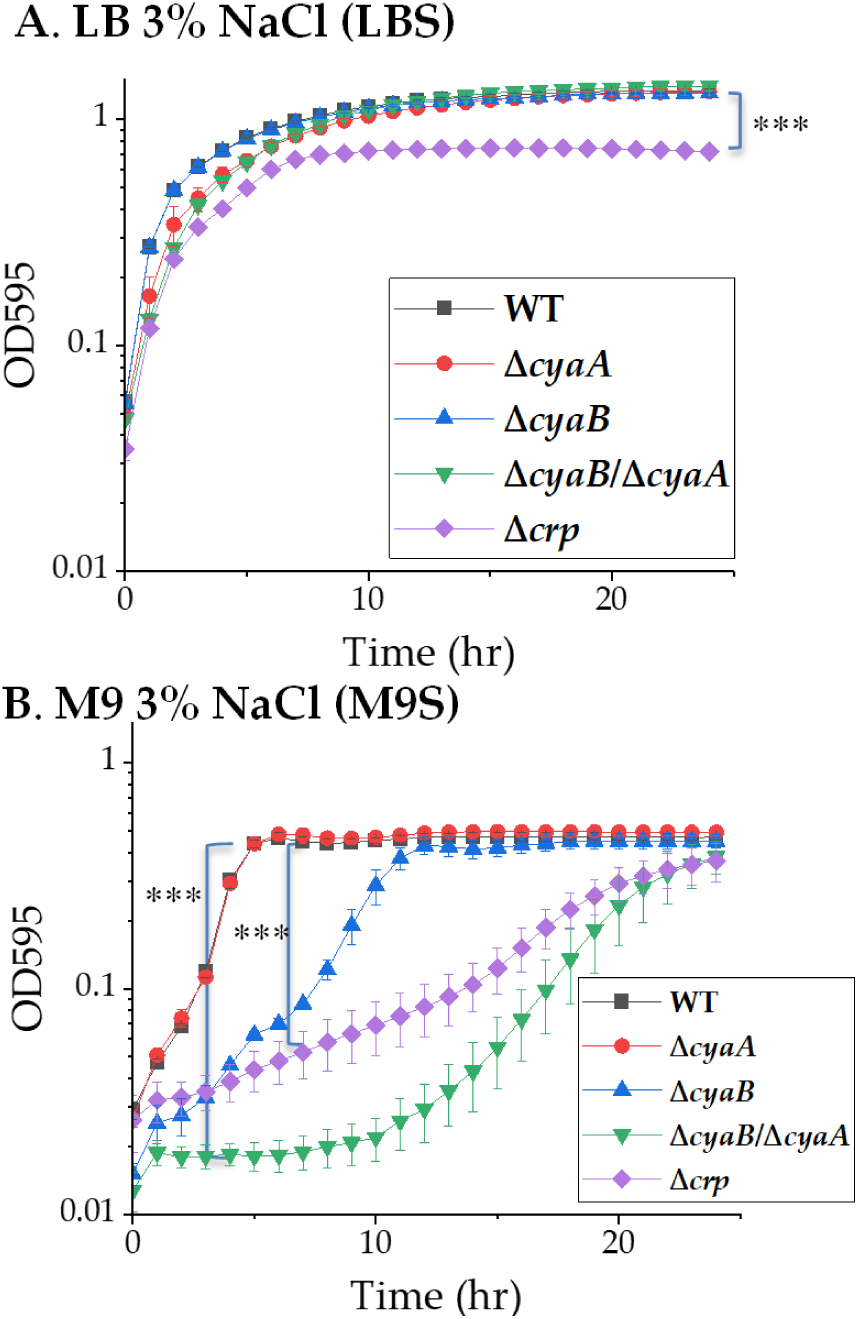
Growth pattern analysis in LBS and M9S D-Glucose. Growth pattern analysis of *V. parahaemolyticus* RIMD2210633 (WT), Δ*cyaA*, Δ*cyaB*, Δ*cyaB/*Δ*cyaA,* and Δ*crp* in **(A)** LBS **(B)** M9S + 10mM D-glucose grown at 37°C with aeration for 24 h. For growth in M9, the strains were grown overnight in M9 3%NaCl (M9S) + 10mM glucose (M9GS) at 37°C with aeration for 24 h. Optical density (OD_595_) was measured every hour for 24 h; means and standard errors of at least two biological replicates are displayed. For all growth curves the area under curve (AUC) was calculated for each strain and a Student’s *t*-test was performed, *P<0.05; *** *P*<0.001.

**Fig. 2.**
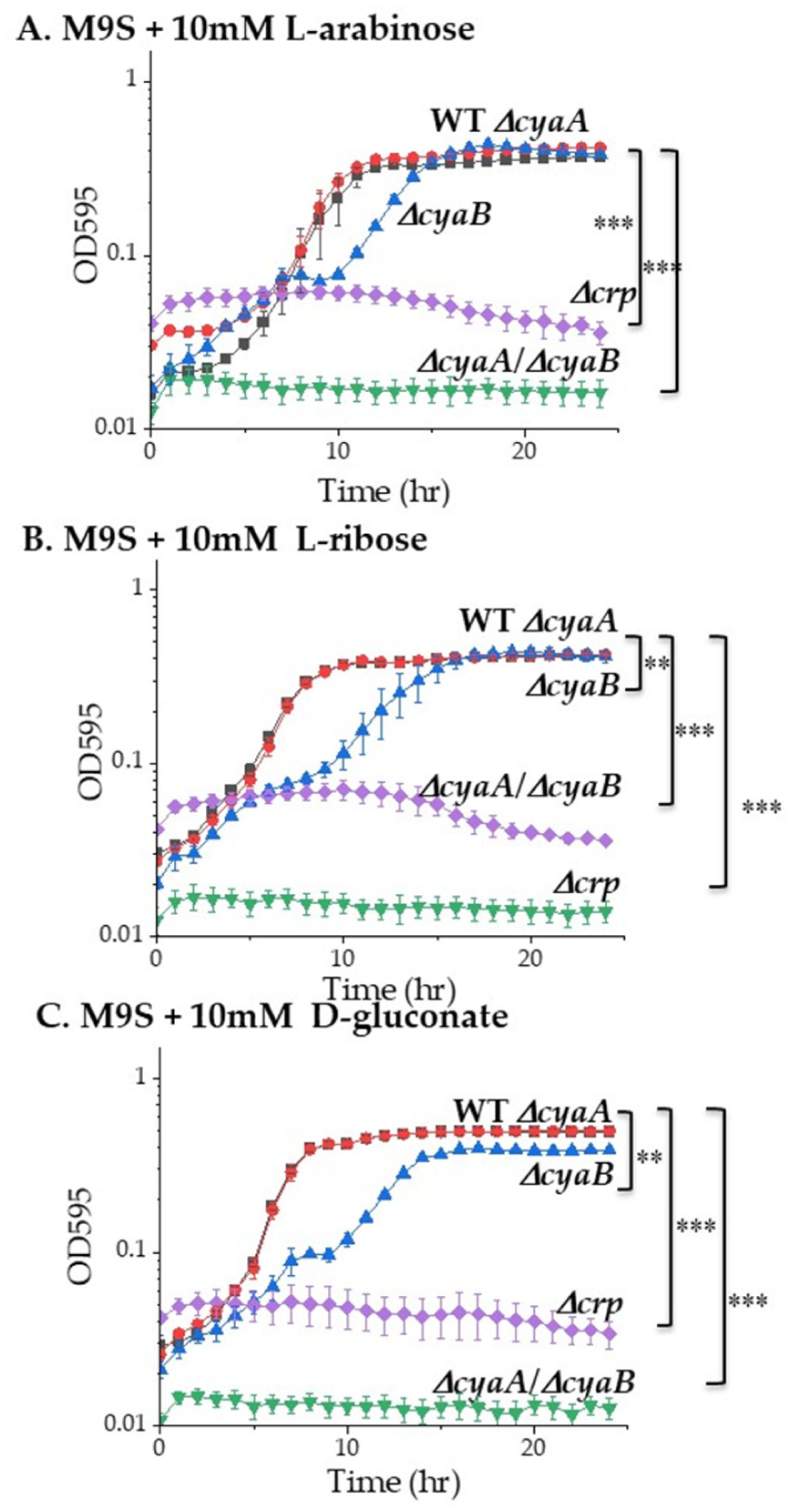
Growth pattern analysis in alternative carbon sources. Growth pattern analysis of *V. parahaemolyticus* RIMD2210633 (WT), Δ*cyaA*, Δ*cyaB*, Δ*cyaB/*Δ*cyaA,* and Δ*crp* in **(A)** M9S + 10mM L-arabinose, **(B)** M9S + 10mM L-ribose **(C)** M9S + 10mM D-gluconate at 37°C with aeration for 24 h. Inoculums were grown overnight in M9S + 10mM glucose at 37°C with aeration. Optical density (OD_595_) was measured every hour for 24 h; means and standard errors of at least two biological replicates are displayed. For all growth curves the area under curve (AUC) was calculated for each strain and a Student’s *t*-test was performed, ** P<0.01; *** *P*<0.001.

### Identification of a class IV adenylate cyclase (AC-IV) CyaB in *V. parahaemolyticus*

BLASTP analysis using CyaA (VP2987) as a seed to search the *V. parahaemolyticus* RIMD2210633 genome, only hit to itself, indicating no other homolog of AC-I was present. Then, a search of the *V. parahaemolyticus* genome for adenylate cyclase classes AC-II using locus WP_080378568 from *Bordetella pertussis*, AC-III using locus PWU33926 from *Pseudomonas aeruginosa*, and AC-IV using locus UNB56521 from *A. hydrophila,* was performed. In addition, ten AC proteins (loci Rv0386, Rv1264, Rv1318c, Rv1319c, Rv1320c, Rv1625c, Rv1647, Rv1900c, Rv2212, Rv3645) encompassing AC-II and AC-III from *Mycobacterium tuberculosis* that were shown to have activity were also used as seeds. These searches identified a homolog of AC-IV from *A. hydrophila* named CyaB. No additional adenylate cyclases were identified in *V. parahaemolyticus* RIMD2210633.

The 181 amino acid CyaB protein lacked transmembrane domains and signal peptide cleavage sites. CyaB contained only one domain and unlike AC-I, AC-II and AC-III does not contain any regulatory or signaling domains. CyaB shared 56% amino acid identity with CyaB of *A. hydrophila* and 45% amino acid identity with CyaB from *Y. pestis*. CyaB from both *A. hydrophila* and *Y. pestis* were shown to have AC activity generating cAMP, however a physiological role was not established (31–34). In *V. parahaemolyticus, cyaB* is encoded on the negative strand upstream of VP1761, which encoded a DUF962-domain hypothetical protein separate by an intergenic region of 88-bp. The AC-IV CyaB protein has been structurally characterized in *Y. pestis* (2FJT and 3NOY) and a crystal structure is also available for VP1760 from *V. parahaemolyticus* (2ACA) (32–34). Comparison of *Y. pestis* and *V. parahaemolyticus* primary sequence and crystal structures shows they are highly similar with active sites conserved between the proteins (**Fig. S3**). The primary sequence of all three bacterial sequences contained the N-terminal conserved HFxxxxExExK motif and the archaea sequences contained the shorter ExExK motif (**Fig. S3**).

### Distribution and phylogenetic analysis of AC-IV among Bacteria

Using *cyaB* (VP1760) as a seed using BLAST analysis, we examined the distribution among nearly 18,000 members of the family *Vibrionaceae* in the NCBI genome database. The *cyaB* gene was present in all characterized *V. parahaemolyticus* strains and in many members of the Harveyi clade, to which *V. parahaemolyticus* belongs, including *V. campbellii, V. harveyi, V. jasicida, V. owensii,* and *V. rotiferanus* (**Fig. 3**). The CyaB proteins in these species showed ∼70% amino acid identity with CyaB from *V. parahaemolyticus,* as compared to ∼95% amino acid identity among the CyaA proteins. On the *cyaB* gene tree, within the Harveyi clade, species formed two divergent branches; one branch representing *V. parahaemolyticus* strains and the second branch containing the remaining species. This suggests that *cyaB* in *V. parahaemolyticus* has diverged significantly from *cyaB* in the other Harveyi clade species. Interestingly, CyaB was absent from most *V. alginolyticus* strains, a species closely related to *V. parahaemolyticus*. Our analysis did identify a *V. alginolyticus* strain that contained a *cyaB* gene on a plasmid that clustered within the same group as *cyaB* present on a plasmid in *V. coralliilyticus* indicating that *cyaB* was acquired by horizontal gene transfer in these species (**Fig. 3**). The *cyaB* gene was present in ∼35 *V. cholerae* strains (from a total of 1,700 genomes in the NCBI database) and two strains of its sister species *V. metoecus* from a total of 30 genomes, which clustered together on the tree. The *cyaB* gene from species within the genera *Aliivibrio* and most *Photobacterium* formed separate divergent branches from *cyaB* in *Vibrio* species (**Fig. 3**). However, *cyaB* from five *Photobacterium* species clustered within *Vibrio* clades suggesting that these *Photobacterium* species acquired a copy of the *cyaB* gene from *Vibrio*, indicating horizontal gene transfer (**Fig. 3**). Comparison of CyaA phylogeny with the CyaB phylogeny from the same set of species also demonstrated additional instances of horizontal transfer among species shown by the non-congruency of the branching patterns for several CyaB proteins (**Fig. S4**).

**Fig. 3.**
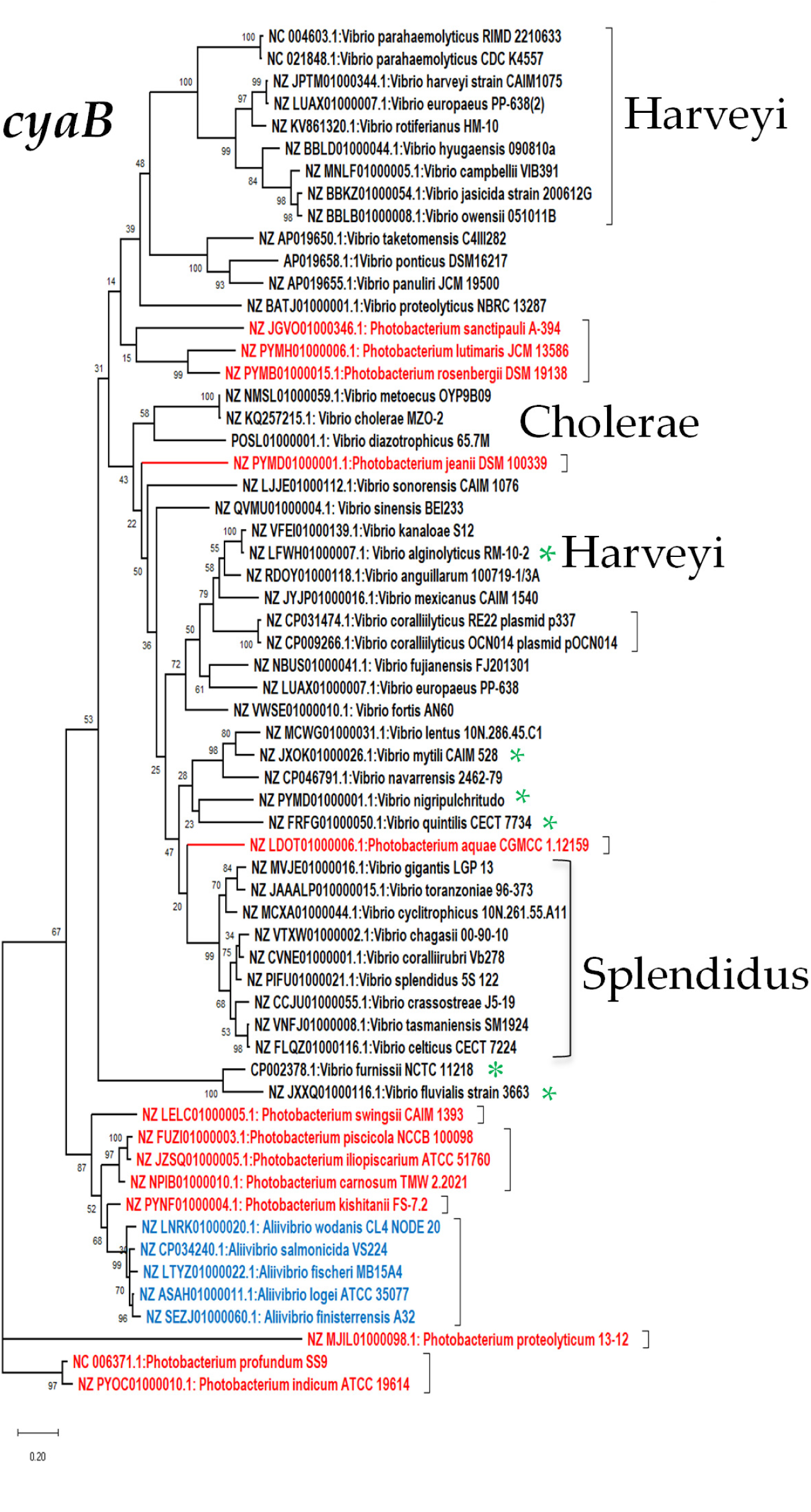
Evolutionary analysis by of *cyaB* among the family *Vibrionaceae*. Phylogenetic analysis was conducted in MEGA X. The evolutionary history was inferred by using the Maximum Likelihood method and Kimura 2-parameter model. The *cyaB* gene tree with the highest log likelihood (-16932.68) is shown. The percentage of trees in which the associated taxa clustered together is shown next to the branches. This analysis involved 61 nucleotide sequences and a total of 543 positions in the final dataset.

Next, we examined the distribution and phylogeny of AC-IV among Bacteria and Archaea genomes in the NCBI genome database (**Fig. 4**). This analysis showed that AC- IV were present predominantly among Gamma-Proteobacteria. CyaB from some species belonging to the genera *Shewanella, Marinomonas, Psychromonas,* and *Morganella* clustered closely with CyaB from *Vibrio* species. Species belonging to the genera *Aeromonas, Yersinia, Brennia, Dickeya, Hafnia, Proteus Providencia,* and *Pectobacterium* contained a copy of CyaB that branched along phylogeny lines indicating that CyaB is ancestral to these lineages (**Fig. 4**). A handful of Alpha-Proteobacteria, *Cohaesibacter, Hoeflea, Kangsaoukella, Maritalea, and Pseudovibrio,* also contained a CyaB homolog that was annotated as AC-IV and these clustered together and branched distantly from CyaB in Gamma-Proteobacteria (**Fig. 4**). Proteins showing greater than 25% amino acid identity with greater than 90% query coverage to CyaB from *V. parahaemolyticus* were present in Archaea and branched divergently from AC-IV homologs in Bacteria (**Fig. 4**). Amino acid sequence comparisons followed by WebLogo analysis of Proteobacteria or Archaea CyaB sequences was performed and generated sequence logos for the entire protein in both groups to identify sequence conservation patterns (**Fig. S5 and S6**). In this analysis, the previously identified AC-IV defining N-terminal motif HFxxxExExK was present in all Gamma-Proteobacteria and Alpha-Proteobacteria sequences examined in this study (**Fig. S5**). In contrast, CyaB representatives from Planctomycete and Archaea contained the N-terminal motif ExExK, which was proposed to be present in CYTH proteins with only phosphatase activity (**Fig. S6**) (35).

**Fig. 4.**
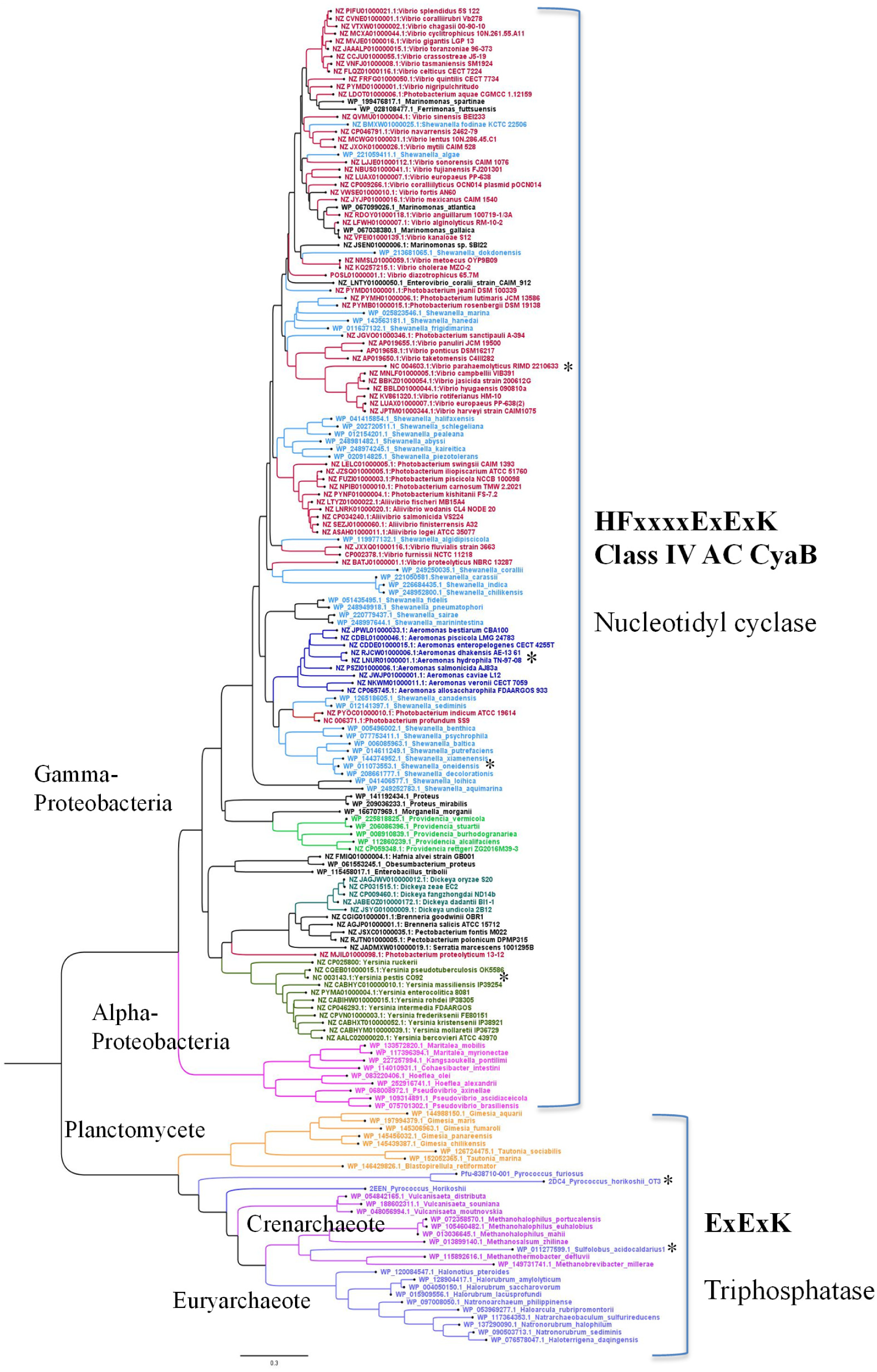
Evolutionary analysis by of CyaB among Gamma- Proteobacteria, Alpha- Proteobacteria, Planctomycete and Archaea species. The CyaB tree was inferred by using the Maximum Likelihood method and Le_Gascuel_2008 model. The tree with the highest log likelihood (-21377.64) is shown. This analysis involved 177 amino acid sequences and a total of 145 positions in the final dataset.

### AC-IV CyaB functionally complements an *E. coli cyaA* deletion mutant

To begin to determine whether CyaB has adenylate cyclase activity, we functionally complemented an *E. coli* Δ*cyaA* mutant with the *cyaA* or *cyaB* gene from *V. parahaemolyticus*. *E. coli* Δ*cyaA*, *E. coli* Δ*cyaA* pBBR*cyaB,* and *E. coli* Δ*cyaA* pBBR*cyaA* were grown in M9S supplemented with glucose as a sole carbon source and all strains grew similarly (**Fig. 5A**). Growth was examined in M9S supplemented with gluconate as a sole carbon source. In this analysis, the growth defect of *E. coli* Δ*cyaA* was rescued in *E. coli* Δ*cyaA* complemented with either pBBR*cyaA* or pBBR*cyaB* (**Fig. 5B)**.

**Fig. 5:**
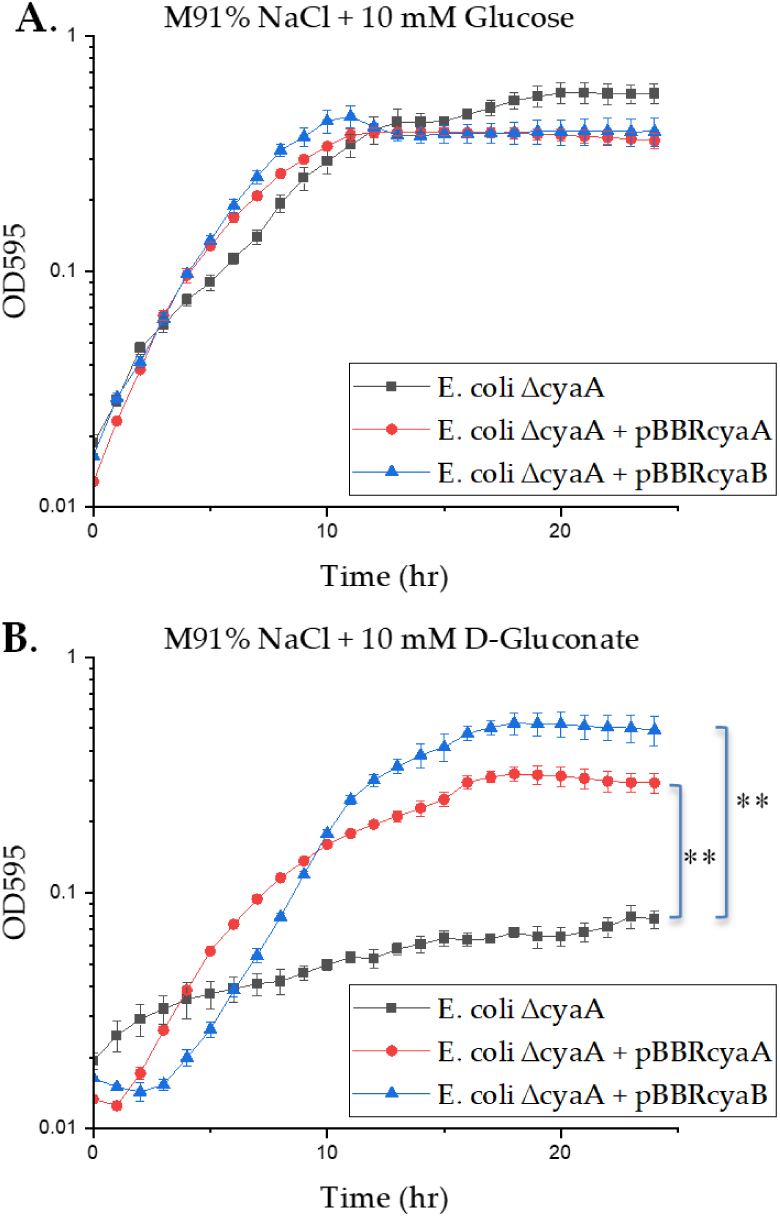
Complementation analysis of *E. coli* Δ*cyaA.* The *E. coli* Δ*cyaA* strain was complemented with a functional copy of *cyaA* or *cyaB* from *V. parahaemolyticus* and growth curves were conducted in **(A)** M9 + 10 mM glucose **(B)** M9 + 10mM gluconate for 48 h at 37°C with aeration. Optical density (OD_595_) was measured every hour for 24 h; means and standard errors of three biological replicates are displayed. For all growth curves the area under curve (AUC) was calculated for each strain and a Student’s *t*-test was performed, ** *P*<0.01.

To investigate the role of *cyaB* in *V. parahaemolyticus* physiology further, we constructed a *cyaB* deletion mutant strain. We examined growth of wild type and the Δ*cyaB* mutant on solid and in liquid media. The Δ*cyaB* mutant had similar colony morphology to wild type on LBS plates (**Fig. S2A**) and showed identical growth patterns in LBS broth to wild type **(Fig. 1A)**. However, when the Δ*cyaB* mutant was grown overnight in M9S D-glucose and used to inoculate M9 supplemented with D-glucose, a longer lag phase than wild type was observed, but a similar biomass was reached **(Fig. 1B)**.

Next, we constructed a Δ*cyaB/*Δ*cyaA* double mutant strain by deleting *cyaA* from the Δ*cyaB* background. The double mutant demonstrated small translucent colony morphology on LBS similar to the colonies observed for Δ*crp* **(Fig. S2A and C)**. Addition of cAMP to the media restored the large colony morphology for the double mutant indicating that the defect was due to the absence of cAMP (**Fig. S2D**).

Interestingly, a glycerol stock of the double mutant when streaked on LBS plates produced small translucent colonies as well as regular opaque colonies, suggesting suppressor mutations were occurring (**Fig. S2B**). We did not observed this phenotype in either of the single Δ*cyaA or* Δ*cyaB* mutants. In *V. cholerae,* which contains only an AC-I

CyaA, the Δ*cyaA* mutant reversion to wild-type phenotypes was also observed (41). For our studies, small colonies were always used for phenotypic assays. The Δ*cyaB/*Δ*cyaA* double mutant showed a growth defect on M9S D-glucose (**Fig. 1B**) and showed no growth in M9 supplemented with L-arabinose, L-ribose, or D-gluconate **(Fig. 2A-C)**.

These growth assays were performed in M9S L-arabinose, L-ribose, or D-gluconate with inoculum obtained after overnight growth in M9S D-glucose. These growth assays showed that the Δ*cyaA* mutant grew similar to wild type, but the Δ*cyaB* mutant growth curve showed a different slope (**Fig. 2**). In contrast, when mutants were grown overnight in LBS and then inoculated into alternative carbon sources, Δ*cyaB* growth curves showed a shorter lag phase and a significantly greater biomass than Δ*cyaA*, which grew similar to wild type (**Fig. S7**). Analysis of the regulatory regions of both AC genes identified putative CRP binding sites in the regulatory region of *cyaA,* but not in the *cyaB* regulatory region (**Fig. S8**). These data suggest that *cyaA* is under CRP control and that *cyaB* is not, and that CyaA and CyaB can complement each other.

### cAMP-CRP regulation of swimming and swarming motility

Unlike *V. cholerae* which only produces a polar flagellum, *V. parahaemolyticus* produces both polar and lateral flagella required for swimming and swarming motility, respectively, which are both activated by the sigma factor RpoN (42). To investigate the role of CRP in *V. parahaemolyticus* motility, we conducted swimming assays and found that the Δ*cyaA* and Δ*cyaB* mutants grew similar to wild type (**Fig. 6A**). The Δ*crp* and Δ*cyaB/*Δ*cyaA* double mutants were defective in swimming compared to wild type, but did not show a complete defect similar to the Δ*rpoN* mutant (**Fig. 6A**). Also, Δ*crp* and Δ*cyaB/*Δ*cyaA* mutant showed a swarming defect as indicated by the absence of the budding phenotype (**Fig. 6B**). The Δ*cyaA* and Δ*cyaB* mutants grew similar to wild type producing typical budding swarming cells as previously shown for this species (**Fig. 6B**) (43–45). The WT, Δ*cyaA* and Δ*cyaB* swarming colonies were examined further to show that the swarming colony structural differences were not due to mutants forming. To accomplish this, different parts of the swarming colony were re-streaked onto swarming plates and examined after 48 h at 30°C, which again showed similar colony morphologies to the original colonies (**Fig. S9**). Since Δ*crp* and Δ*cyaB/cyaA* both gave a small colony morphology on swarming plates after 48 h of growth at 30°C, we extended the incubation time to 64 h at 30°C to ensure no swarming motility occurred **(Fig. S10).** Overall, the data demonstrate that cAMP-CRP is an activator of both swimming and swarming motility in *V. parahaemolyticus*.

**Fig. 6.**
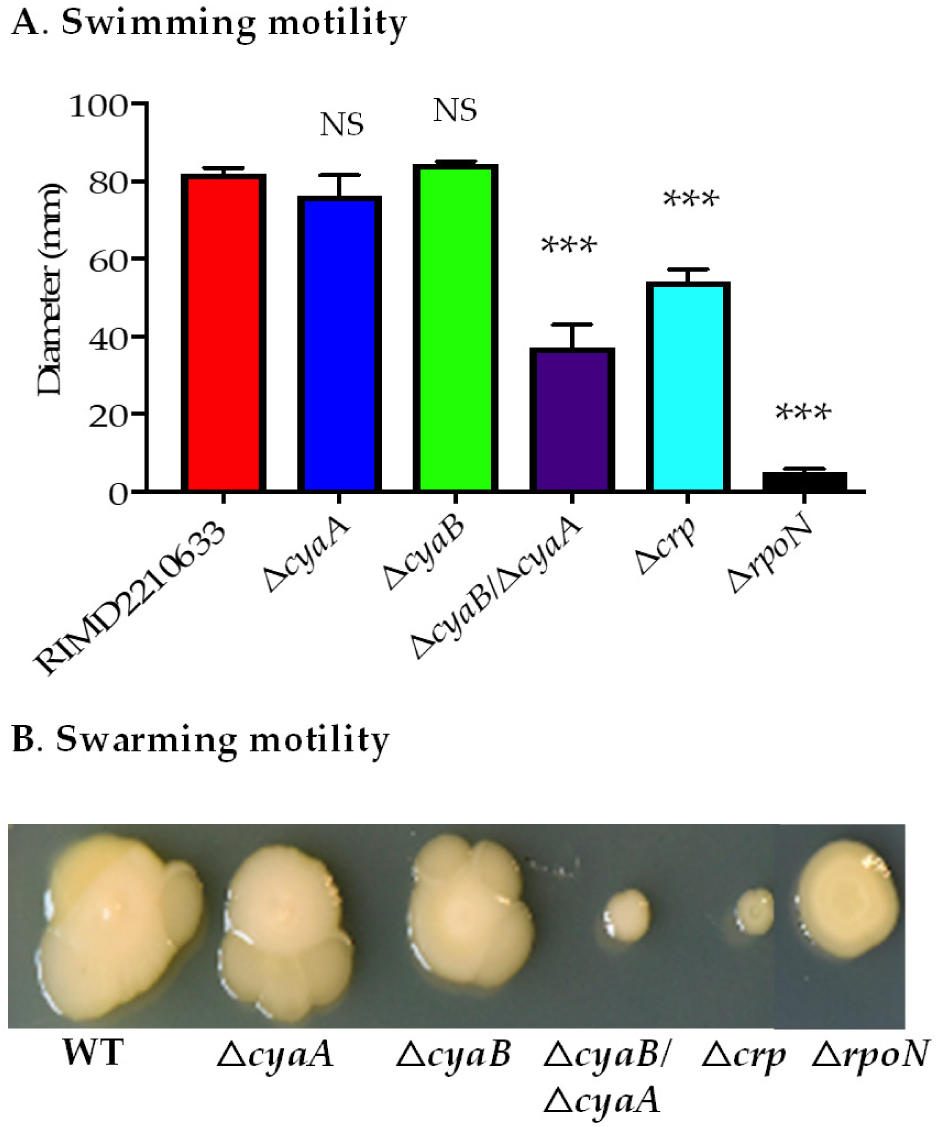
Characterization of motility in *V. parahaemolyticus* mutants. (A) Swimming assays were conducted using wild type, Δ*crp,* Δ*cyaA,* Δ*cyaB*, and Δ*cyaB/*Δ*cyaA* mutants grown at 37°C for 24 h in LB 2% NaCl with 0.3% agar. The diameter of the region covered by the bacterial strains were measured and represented as bar graph. The error bars indicate standard error of mean and an Unpaired Student t-test was conducted. The significant difference is denoted by asterisks ***, P<0.001 and NS indicates not significant **(B**) Swarming assays. Colonies of wild type and mutant strains Δ*crp,* Δ*cyaA,* Δ*cyaB*, and Δ*cyaB/*Δ*cyaA* were assessed for swarming motility in HI containing 2% NaCl and 1.5% agar at 30°C for 48 h.

### cAMP-CRP and AC-IV CyaB are required for capsule and biofilm production

Capsule polysaccharide (CPS) is an important component of biofilm and is positively controlled by the quorum sensing master regulator OpaR in *V. parahaemolyticus* (43, 44, 46–49). To determine whether CRP plays a role in CPS formation, we examined the Δ*crp*, Δ*cyaA,* Δ*cyaB,* and Δ*cyaB/cyaA* mutants on HI medium containing Congo red dye and CaCl_2._ After incubation for 48 h at 30°C, the wild type strain produced a rugose opaque colony as did the Δ*cyaA* mutant, both indicatingf CPS production (**Fig. 7**). However, the Δ*cyaB* mutant produced a smooth translucent colony as did the Δ*crp* and Δ*cyaB*/Δ*cyaA* mutants indicating absence of CPS (**Fig. 7**). In biofilm assays, wild type and Δ*cyaA* produced significant biofilm, whereas Δ*crp*, Δ*cyaB,* and Δ*cyaB*/Δ*cyaA* produced significantly less biofilm compared to wild type (**Fig. 8**). Overall, these data demonstrate that both CRP and CyaB, but not CyaA, are required for CPS and biofilm production in *V. parahaemolyticus*. We functionally complemented a *V. parahaemolyticus* Δ*cyaB* mutant with the *cyaB* gene on an expression vector and a biofilm assay was performed. In this assay the wild type pBBREV (empty vector), Δ*cyaB* pBBREV, and Δ*cyaB* pBBR*cyaB* were examined and the Δ*cyaB* pBBR*cyaB* complement showed restoration of biofilm to wild type levels (**Fig. S11**).

**Fig. 7.**
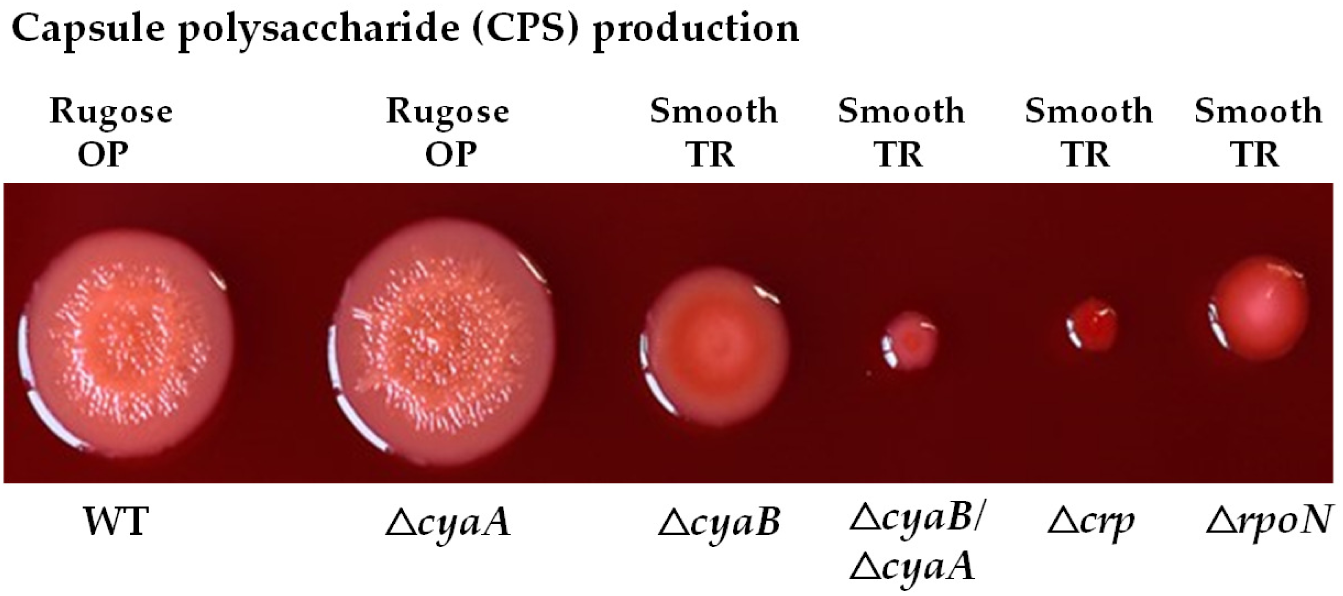
Capsule polysaccharide production. Colonies of *V. parahaemolyticus* RIMD2210633 and mutant strains were inoculated on HI plate containing Congo red dye and CaCl_2_ grown at 37°C for 24 h.

**Fig. 8.**
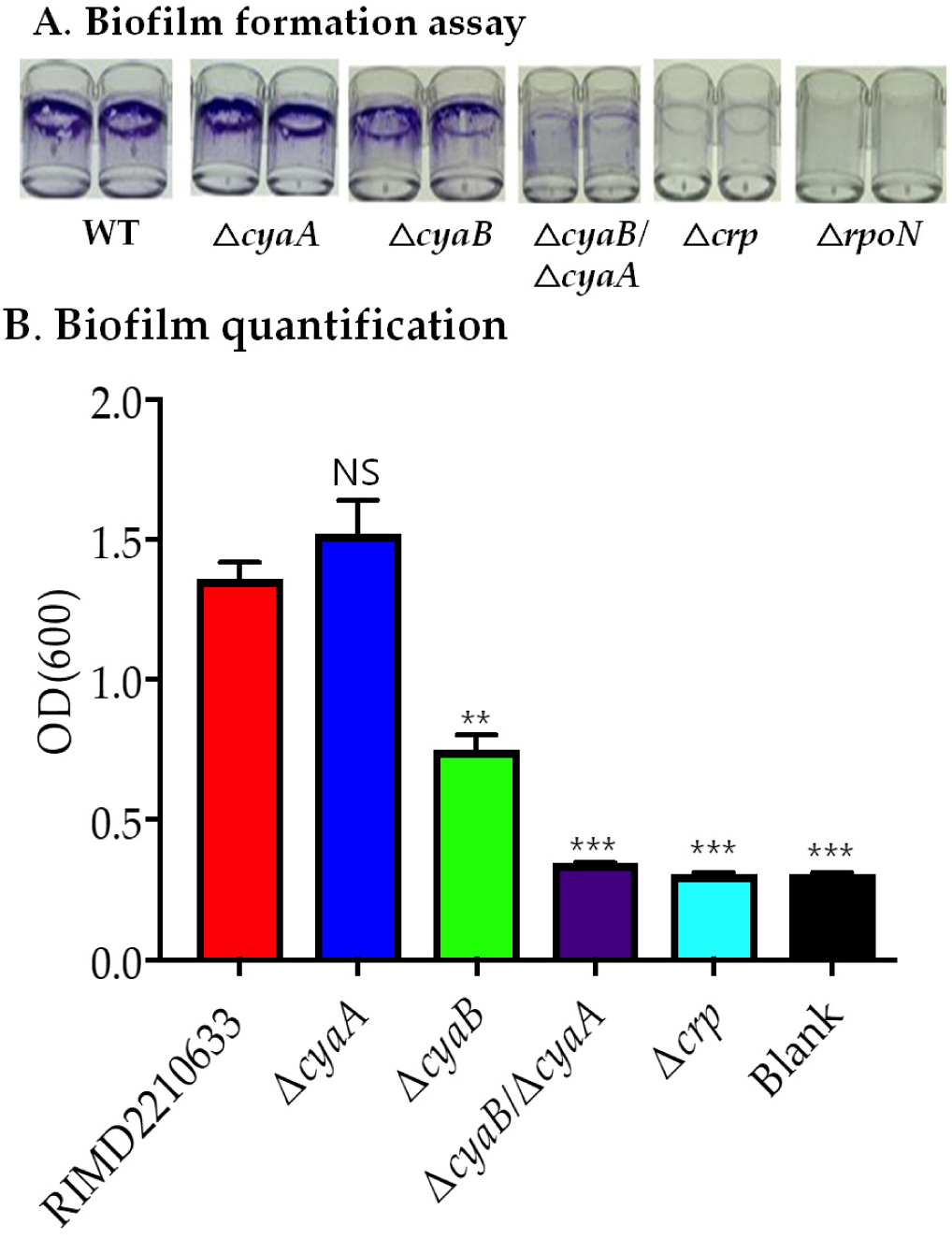
Biofilm formation. *V. parahaemolyticus* strains were grown in HI broth for 24 h. **A**. The biofilms were visualized using crystal violet and then dissolved using DMSO, and the optical density of the biofilms were measured at OD and **B.** represented as bar graphs. Error bars indicate standard error of mean (SEM). Unpaired Student *t* test was conducted to obtain the *P* values. The significant difference in biofilm production is denoted by asterisks. *, P<0.05, **, P<0.01, ***, P<0.001. NS, not significant.

Both Δ*crp* and Δ*opaR* deletion mutants have significant defects in capsule formation and biofilm production. It was previously shown that OpaR is an activator of CPS binding to the regulatory regions of the CpsA-K biosynthesis cluster and CPS regulators CspQ, CpsR, and CpsS in *V. parahaemolyticus* BB22OP (50). We examined whether CRP and OpaR interact to control CPS production. To accomplish this, functional complement experiments of Δ*crp* and Δ*opaR* with either *crp* or *opaR* were performed. As controls, both Δ*crp* and Δ*opaR* were functionally complemented with their respective gene. The Δ*opaR* pBAD*opaR* showed restoration of CPS and Δ*crp* pBAD*crp* produced a large non- translucent colony (**Fig. 9A**). The Δ*opaR* pBAD*crp* strain remained translucent, suggesting that *crp* cannot rescue the Δ*opaR* CPS defect (**Fig. 9A**). Also, the Δ*crp* pABAD*opaR* mutant did not appear to restore CPS (**Fig. 9A**), however in biofilm assays, Δ*crp* pBAD*crp* produced significantly more biofilm than Δ*crp* pBADEV (**Fig. 9B, Fig. 9C**). In the Δ*crp* pBAD*opaR* strain, significantly more biofilm was produced compared to Δ*crp* pBADEV, but not to the same level as Δ*crp* pBAD*crp* **(Fig. 9D)**. These data suggest that CRP is epistatic to OpaR in the biofilm regulatory network, but additional factors are involved in biofilm regulation, which needs to be examined further.

**Fig. 9.**
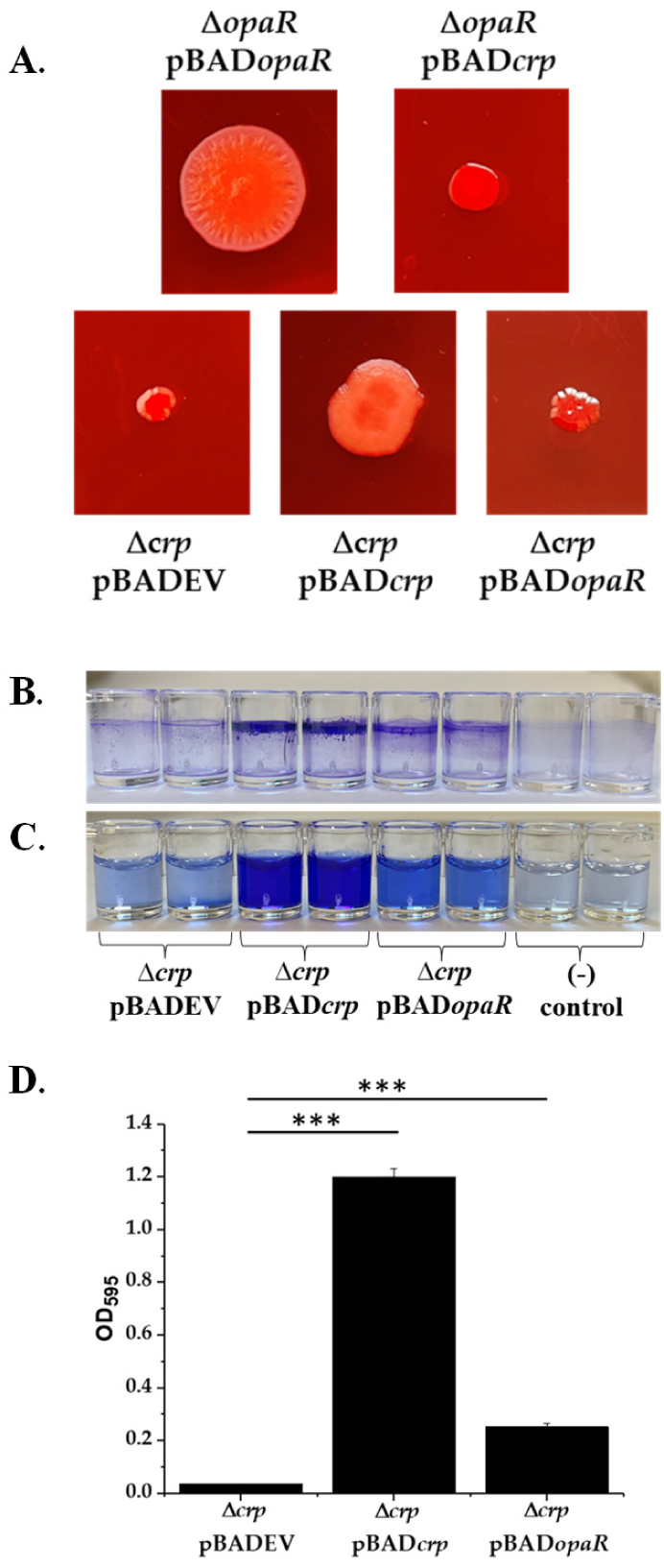
**A**. Capsule polysaccharide (CPS) assays. The pBAD*opaR* and pBAD*crp* complementation plasmids were conjugated into *V. parahaemolyticus* mutant strains. Cells were grown on HI congo red CaCl_2_ media with 0.1% L-arabinose and 12.5 μg/mL chloramphenicol at 30°C for 48 h prior to imaging. **B.** Biofilm production assays. *V. parahaemolyticus* Δ*crp* mutant containing pBAD33 (pBADEV), pBAD*crp*, or pBAD*opaR* expression plasmids were grown in 3% LB with 12.5 μg/mL chloramphenicol and 0.1% L-arabinose for 24 hours at 37°C. **C.** Biofilm solubilized in DMSO. **D**. Biofilm measured for absorbance at OD_595_. Error bars indicate standard error of mean (SEM). An Unpaired Student *t* test was conducted to obtain ***, P<0.001.

## DISCUSSION

In this study, the physiological roles of CRP and adenylate cyclase were examined in *V. parahaemolyticus*, which led to the identification of an uncharacterized AC-IV named CyaB. The data showed that in *V. parahaemolyticus,* CRP was an important regulator of metabolism and motility, acting as a positive regulator of both swimming and swarming motility. In addition, the data showed that CRP was required for capsule and biofilm formation. In *E. coli* and other enterobacteria deleting *cyaA,* which encodes the canonical AC-I, results in CRP remaining inactive. Thus in these species, the Δ*cyaA* and Δ*crp* mutants have similar phenotypes (51, 52). In *V. cholerae* and *V. vulnificus,* deletion of *cyaA* also gave phenotypes similar to Δ*crp,* showing defects in metabolism and motility (38, 53, 54). In *V. parahaemolyticus,* the Δ*cyaA* mutant showed phenotypes similar to wild type. This unexpected finding led us to search for proteins with putative AC function in the *V. parahaemolyticus* genome leading to the identification of the *cyaB* gene (VP1760). CyaB belongs to the CYTH protein superfamily (Pfam PF01928) and was annotated as a class AC-IV, which contained one domain and was homologous to AC- IV proteins previously described in *A. hydrophila*, *Y. pestis,* and thermophilic archaebacteria (31–34). Our phylogenetic analysis showed that CyaB is widespread throughout the *Vibrionaceae* family generally present in all strains of each species.

Several instances of horizontal gene transfer of *cyaB* were noted. Overall, AC-IV homologs were present predominantly among Gamma-Proteobacteria in species belonging to the genera *Aeromonas, Shewanella, Yersinia, Proteus, Providencia, Dickeya,* and *Pectobacterium,* amongst others. In these genera, the CyaB phylogeny usually followed evolutionary relationships, but some *Shewanella* species for example nested within *Vibrio* CyaB clades suggesting horizontal gene transfer. Homologs were also present in Alpha-Proteobacteria, but only in a handful of genera; *Cohaesibacter, Hoeflea, Kangsaoukella, Maritalea, and Pseudovibrio.* Outside of Bacteria, proteins annotated as AC- IV were present in Planctomycete, Crenarchaetote and Euryarchaeote species. All bacterial homologs of AC-IV described in this study contained the N-terminal motif HFxxxxExExK. The amino acid residues HF are an important distinguishing feature present in proteins with nucleotidyl cyclase activity, whereas among Planctomycete and Archaea species only the N-terminal motif ExExK is present. Vogt and colleagues recent work showed that archaeal CYTH-like proteins are functionally diverge from bacterial AC-IV CyaB-like proteins (35). They proposed that a true AC-IV requires the extended N-terminal signature HFxxxxExExK, with the HF motif absent in other CYTH proteins that function as either TTM or ThTPase (35).

The seminal studies of AC-IV in *A. hydrophila* showed that the protein was only active at high temperatures and alkaline pH conditions (31). Furthermore, their studies showed that deleting *cyaB* did not have any cellular effects under the conditions examined (31). A study in *Shewanella oneidensis* characterized three AC genes, *cyaA, cyaB,* and *cyaC* members of AC-I, AC-IV, and AC-III present in that species. They showed each *cya* gene functionally complemented an *E. coli* Δ*cyaA*, but did not determine any other role for *cyaB* (55). Indeed, to date, no physiological role for AC-IV has been shown. In our study, we show that deleting *cyaB* results in significant defects in metabolism, capsule production, and biofilm formation. The defects in CPS production and biofilm formation in both the Δ*crp* and Δ*cyaB* mutants indicate that cAMP-CRP is a positive regulator of these phenotypes and CyaB is essential for this function. The data may also suggest that cAMP, per se, is not directly involved, but that a function specific to the CYTH superfamily could be.

Studies in *V. cholerae* identified cAMP-CRP as a negative regulator of capsule production and biofilm formation (38, 56). It was showed that cAMP-CRP negatively regulated CPS (encoded by *vsp* loci) and biofilm matrix proteins by regulating genes encoding diguanylate cyclases and phosphodiesterases. In *V. cholerae*, the *vsp* biosynthetic operon is negatively controlled by HapR, the quorum sensing master regulator, and it was shown that CRP was epistatic to HapR in the regulation of the VspT and VspR regulators of capsule production (38, 56–59). In these studies both Δ*crp* and Δ*cyaA* showed decreased expression of *hapR* suggesting that cAMP-CRP was a positive regulator of HapR (38, 56, 60). In *V. parahaemolyticus*, the quorum sensing master regulator OpaR is a positive regulator of capsule and biofilm production, however, whether cAMP-CRP controls OpaR is unknown (43, 44, 46–50). Our preliminary bioinformatics analysis identified two putative CRP binding sites in the regulator region of OpaR suggesting CRP may be a direct regulator of *opaR* in *V. parahaemolyticus* (**Fig. S12**). Putative CRP binding sites were also identified in *cspA-K* regulator promoter regions, which will need to be investigated further (**Fig. S12**). It is of interest to note that although both *V. cholerae* and *V. parahaemolyticus* use conserved regulatory pathways, QS and CCR, to modulate capsule and biofilm production, the effects are the direct opposite in each species. It will be of interest to determine how this diversity in the control of important phenotypes in these related species has evolved and how CyaB plays a role.

## MATERIALS AND METHODS

### Bacterial strains, media, and culture conditions

All strains and plasmids used in this study are listed in **Table 1**. A streptomycin-resistant clinical isolate *V. parahaemolyticus* RIMD2210633 was used throughout the study. Unless stated otherwise, all *V. parahaemolyticus* strains were grown in Luria Bertani (LB) medium (Fischer Scientific, Pittsburgh, PA) containing 3% NaCl (LBS) at 37°C with aeration or M9 medium media (Sigma Aldrich, St. Louis, MO) containing 3% NaCl (M9S) supplement with 10mM of various carbon sources. Antibiotics were added to growth media at the following concentrations: Ampicillin (Amp), 100 μg/ml, streptomycin (Sm), 200 μg/ml, chloramphenicol (Cm), 10 μg/ml, and kanamycin (Km), 50 μg/ml when required.

**Table 1.**
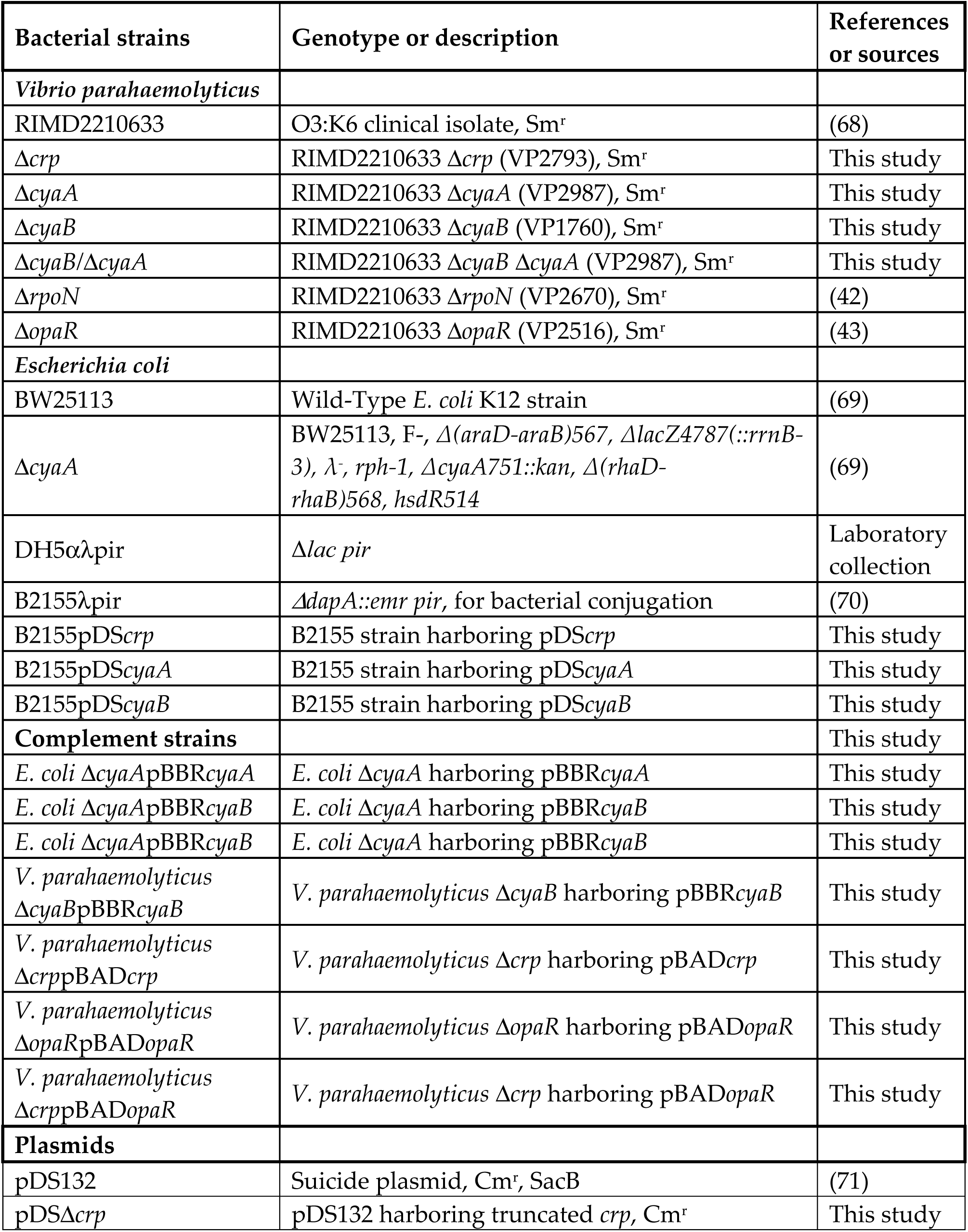
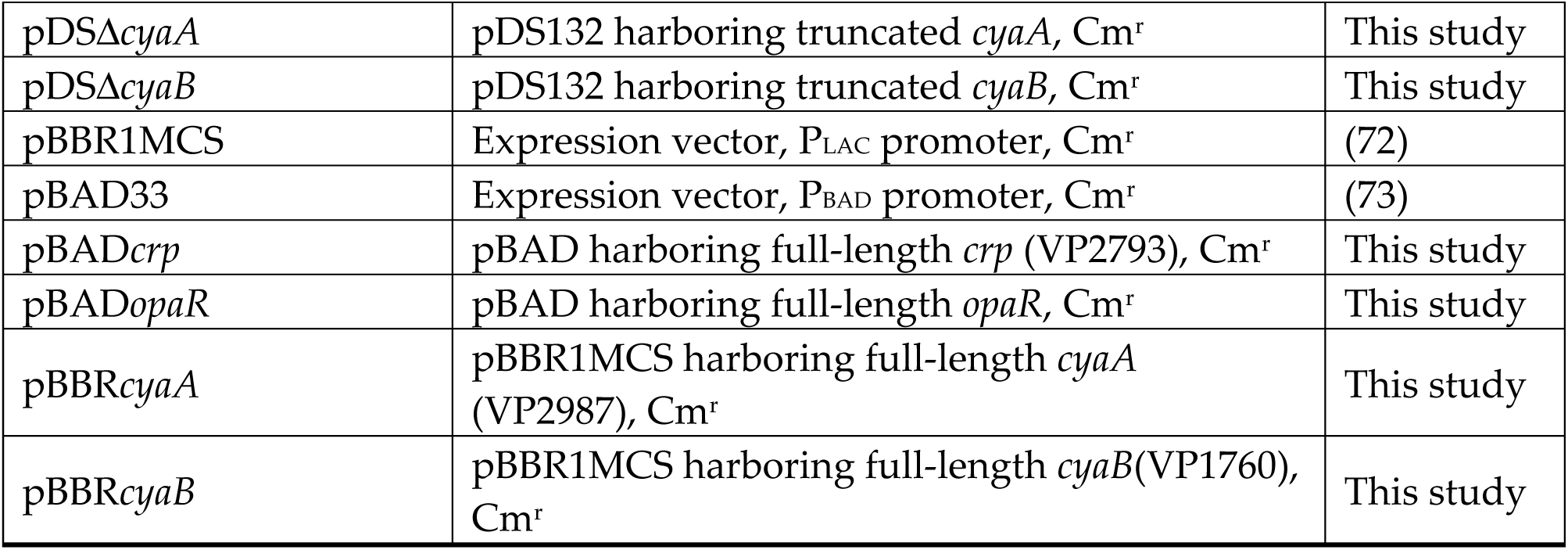
Bacterial strains and plasmids used in this study.

### Construction of mutant strains in *V. parahaemolyticus* RIMD2210633

Splice by overlapping extension (SOE) PCR and an allelic exchange (61) was used to construct in-frame, non-polar deletion mutants of *crp* (VP2793), *cyaA* (VP2987), *cyaB* (VP1760) and a double deletion *cyaB/cyaA* in *V. parahaemolyticus* RIMD2210633. Primer pairs were designed for *crp* (VP2793), *cyaA* (VP2987), and *cyaB* (VP1760) PCR amplification using *V. parahaemolyticus* RIMD2210633 genomic DNA as a template. All primers used in this study are listed in **Table S1**. Using the PCR amplified fragments of each gene, SOE PCR was conducted to create truncated version of *crp* (leaving 81-bps of 633-bp VP2793), *cyaA* (153-bp of 2,529-bp VP2987) and *cyaB* (84-bp of 546-bp VP1760).

The truncated PCR fragments were cloned into the suicide vector pDS132 and named pDS*Δcrp,* pDS*ΔcyaA,* and pDS*ΔcyaB*. pDS*Δcrp,* pDS*ΔcyaA,* and pDS*ΔcyaB* were transformed into *E. coli* strain β2155 λ*pir*, which were conjugated separately into *V. parahaemolyticus*. Conjugation was conducted by spread plating both strains onto LBS plate containing 0.3 mM diaminopimelic acid (DAP). Bacterial growth from this plate was scraped, suspended in LBS, serially diluted, and plated on LBS plates containing streptomycin and chloramphenicol. After overnight growth, colonies were examined to identify single crossovers of the suicide plasmids into the target site via PCR. The colonies with single crossover were grown over night in LBS with no antibiotic and plated onto LBS plate containing 10% sucrose to select for double crossover deletion mutants. The double deletion mutant Δ*cyaB/*Δ*cyaA* was constructed by deleting *cyaA* from Δ*cyaB* background following the above protocol. The gene deletions were confirmed by PCR and DNA sequencing.

### Bioinformatics analysis

The presence of CRP binding sites (BS) in the regulatory regions of *crp, cyaA* and *cyaB* in *V. parahaemolyticus* were examined using PRODORIC software using both the *E. coli* and *V. cholerae* consensus sequences for Crp (62). WebLogo was used to generate graphical representations of the multiple sequence alignments used to generate phylogenetic trees for CyaB (63). Each single letter symbol is the single letter designation of an amino acid at each position in the protein. The height of the each stack for each position indicates the sequence conservation at that position and the height of each symbol within a stack represent the frequency of each amino acid at that position.

### Functional complementation of *E. coli* Δ*cyaA* and *V. parahaemolyticus* mutants

Primer pairs were designed to amplify *cyaA* (VP2987) and *cyaB* (VP1760) using wild type genomic DNA as template (**Table S1**). Purified PCR products of *cyaA* or *cyaB*, and the vector pBBR1MCS were digested using BamHI and XbaI. Both the insert and the vector were gel purified. The genes were each cloned into the vector pBBR1MCS creating pBBR*cyaA* or pBBR*cyaB*, which were then transformed into *E. coli* B2155 using the calcium chloride transformation method. *E. coli* β2155 containing pBBR*cyaA* and pBBR*cyaB* were transformed into *E. coli* Δ*cyaA* and grown overnight in media supplemented with 100 mM isopropyl-β-d-thiogalactopyranoside (IPTG) (64). For functional complementation of *V. parahaemolyticus* Δ*crp,* Δ*cyaB* and Δ*opaR* a similar approach was used, but *crp* and *opaR* as well as *cyaB* were cloned into pBAD33. Complement strains were verified using PCR and sequencing.

### Growth pattern analysis

Bacterial cultures were grown overnight in LBS or M9S supplemented with 10mM glucose at 37°C with aeration. An overnight growth culture was spun down, resuspended, and washed with PBS and a 1: 40 dilution was then used to conduct growth pattern analysis in M9S supplemented with 10 mM of each individual carbon sources. Growth curves were performed in a 96 well plate at 37°C with aeration. Optical density (OD595) was measured each hour for 24 hours. Each experiment was performed in triplicate with two biological replications.

### Swimming and swarming motility assay

Swimming assays were performed in semi-solid media, LB 2% NaCl with 0.3% agar and swarming assays were performed in heart infusion (HI) medium (Remel, Lenexa, KS) containing 2% NaCl and 1.5 % agar. To examine swimming behavior, a single colony of the test bacterium was picked and stabbed in the center of the LB plate and incubated at 37°C for 24 h. For the swarming assay, a single colony was picked and gently inoculated on the surface of the HI plate, and the plates were incubated at 30°C for 48 h. Each experiment was performed in triplicate with three biological replications.

### Capsule polysaccharide and biofilm production assay

Capsule polysaccharide (CPS) formation assays were conducted as previously described (43–45). Wild type and mutant strains were grown on HI media containing 1.5% agar, 2% NaCl, 2.5 mM CaCl_2_, and 0.25% Congo red dye for 48 h at 30°C. Each image is an example from at least three biological replicates performed in triplicate. Biofilm assays were performed as previously described (43–45). Briefly, overnight cultures were used to inoculate strip wells in a 1:40 dilution in LBS. Cultures were incubated for 24 h at 37°C prior to analysis. Biofilm was stained with crystal violet and solubilized with dimethyl sulfoxide (DMSO) (ThermoFisher Scientific, USA). Optical density was measured at 595nm to quantify biofilm.

### Phylogenetic analysis of AC-IV

pBLAST searches with VP1760 from *V. parahaemolyticus* used as a seed were performed on all completed and draft genomes published on the National Center for Biotechnology Institute (NCBI) website. Sequences were aligned using CLUSTALW (65). The evolutionary history was inferred by using the Maximum Likelihood method and Kimura 2-parameter model (66). The *cyaB* gene tree with the highest log likelihood (-16932.68) is shown. The percentage of trees in which the associated taxa clustered together is shown next to the branches. A discrete Gamma distribution was used to model evolutionary rate differences among sites (3 categories (+*G*, parameter = 0.5711)). This analysis involved 61 nucleotide sequences. Codon positions included were 1st+2nd+3rd. There were a total of 543 positions in the final dataset. The CyaB tree was inferred by using the Maximum Likelihood method and Le_Gascuel_2008 model. The tree with the highest log likelihood (-21377.64) is shown. A discrete Gamma distribution was used to model evolutionary rate differences among sites (3 categories (+*G*, parameter = 0.7213)). This analysis involved 177 amino acid sequences and a total of 145 positions in the final dataset. Initial tree(s) for the heuristic search were obtained automatically by applying Neighbor-Join and BioNJ algorithms to a matrix of pairwise distances estimated, and then selecting the topology with superior log likelihood value. Trees are drawn to scale, with branch lengths measured in the number of substitutions per site. All positions with less than 95% site coverage were eliminated, i.e., fewer than 5% alignment gaps, missing data, and ambiguous bases were allowed at any position (partial deletion option). Phylogenetic analyses were conducted with MEGA X (67). For the *Vibrio* species that contained CyaB, a second tree was constructed for the same set of species based on CyaA protein sequences and compared with the CyaA tree to examine differences in branching patterns amongst species.

## ACKNOWLEDGMENTS

This research was supported in part by a National Science Foundation grant (award IOS-1656688) to E.F.B. J.G.T was funded by a University of Delaware graduate fellowship award.

## Notes

### Competing Interest Statement

The authors have declared no competing interest.

### Summary of Updates

New figure 4, new figure 9 and new supplementary data

## REFERENCES

1. Botsford JL, Harman JG. 1992. Cyclic AMP in prokaryotes. Microbiol Rev 56:100–22.

2. Görke B, Stülke J. 2008. Carbon catabolite repression in bacteria: many ways to make the most out of nutrients. Nature Reviews Microbiology 6:613–624.

3. Deutscher J. 2008. The mechanisms of carbon catabolite repression in bacteria. Curr Opin Microbiol 11:87–93.

4. Deutscher J, Francke C, Postma PW. 2006. How phosphotransferase system- related protein phosphorylation regulates carbohydrate metabolism in bacteria. Microbiol Mol Biol Rev 70:939–1031.

5. Danchin A, Guiso N, Roy A, Ullmann A. 1984. Identification of the *Escherichia coli* cya gene product as authentic adenylate cyclase. J Mol Biol 175:403–8.

6. Dumay V, Danchin A, Crasnier M. 1996. Regulation of *Escherichia coli* adenylate cyclase activity during hexose phosphate transport. Microbiology (Reading) 142 (Pt 3):575–583.

7. Makman RS, Sutherland EW. 1965. Adenosine 3’,5’-Phosphate in *Escherichia coli*. J Biol Chem 240:1309–14.

8. Stülke J, Hillen W. 1999. Carbon catabolite repression in bacteria. Curr Opin Microbiol 2:195–201.

9. Kolb A, Busby S, Buc H, Garges S, Adhya S. 1993. Transcriptional regulation by cAMP and its receptor protein. Annu Rev Biochem 62:749–95.

10. Zheng D, Constantinidou C, Hobman JL, Minchin SD. 2004. Identification of the CRP regulon using in vitro and in vivo transcriptional profiling. Nucleic Acids Res 32:5874–93.

11. Roy A, Danchin A. 1982. The cya locus of *Escherichia coli* K12: organization and gene products. Mol Gen Genet 188:465–71.

12. Roy A, Danchin A, Joseph E, Ullmann A. 1983. Two functional domains in adenylate cyclase of *Escherichia coli*. J Mol Biol 165:197–202.

13. Berg OG, von Hippel PH. 1988. Selection of DNA binding sites by regulatory proteins. II. The binding specificity of cyclic AMP receptor protein to recognition sites. J Mol Biol 200:709–23.

14. Benoff B, Yang H, Lawson CL, Parkinson G, Liu J, Blatter E, Ebright YW, Berman HM, Ebright RH. 2002. Structural basis of transcription activation: the CAP-alpha CTD-DNA complex. Science 297:1562–6.

15. Nelson SO, Wright JK, Postma PW. 1983. The mechanism of inducer exclusion. Direct interaction between purified III of the phosphoenolpyruvate:sugar phosphotransferase system and the lactose carrier of *Escherichia coli*. EMBO J 2:715–20.

16. Inada T, Kimata K, Aiba H. 1996. Mechanism responsible for glucose-lactose diauxie in *Escherichia coli*: challenge to the cAMP model. Genes Cells 1:293–301.

17. Hogema BM, Arents JC, Bader R, Eijkemans K, Yoshida H, Takahashi H, Aiba H, Postma PW. 1998. Inducer exclusion in *Escherichia coli* by non-PTS substrates: the role of the PEP to pyruvate ratio in determining the phosphorylation state of enzyme IIAGlc. Mol Microbiol 30:487–98.

18. Saier MH, Jr., Feucht BU, Hofstadter LJ. 1976. Regulation of carbohydrate uptake and adenylate cyclase activity mediated by the enzymes II of the phosphoenolpyruvate: sugar phosphotransferase system in *Escherichia coli*. J Biol Chem 251:883–92.

19. Ullmann A, Tillier F, Monod J. 1976. Catabolite modulator factor: a possible mediator of catabolite repression in bacteria. Proc Natl Acad Sci U S A 73:3476–9.

20. Danchin A, Dondon L, Daniel J. 1984. Metabolic alterations mediated by 2- ketobutyrate in *Escherichia coli* K12. Mol Gen Genetics 193:473–478.

21. You C, Okano H, Hui S, Zhang Z, Kim M, Gunderson CW, Wang YP, Lenz P, Yan D, Hwa T. 2013. Coordination of bacterial proteome with metabolism by cyclic AMP signalling. Nature 500:301–6.

22. Bârzu O, Danchin A. 1994. Adenylyl cyclases: a heterogeneous class of ATP- utilizing enzymes. Prog Nucleic Acid Res Mol Biol 49:241–83.

23. Ramakrishnan G, Jain A, Chandra N, Srinivasan N. 2016. Computational recognition and analysis of hitherto uncharacterized nucleotide cyclase-like proteins in bacteria. Biol Direct 11:27.

24. Crasnier M, Dumay V, Danchin A. 1994. The catalytic domain of *Escherichia coli* K-12 adenylate cyclase as revealed by deletion analysis of the cya gene. Mol Gen Genet 243:409–16.

25. Ladant D, Ullmann A. 1999. Bordatella pertussis adenylate cyclase: a toxin with multiple talents. Trends Microbiol 7:172–6.

26. Sinha SC, Sprang SR. 2006. Structures, mechanism, regulation and evolution of class III nucleotidyl cyclases. Rev Physiol Biochem Pharmacol 157:105–40.

27. Cotta MA, Whitehead TR, Wheeler MB. 1998. Identification of a novel adenylate cyclase in the ruminal anaerobe, Prevotella ruminicola D31d. FEMS Microbiol Lett 164:257–60.

28. Téllez-Sosa J, Soberón N, Vega-Segura A, Torres-Márquez ME, Cevallos MA. 2002. The Rhizobium etli cyaC product: characterization of a novel adenylate cyclase class. J Bacteriol 184:3560–8.

29. Iyer LM, Aravind L. 2002. The catalytic domains of thiamine triphosphatase and CyaB-like adenylyl cyclase define a novel superfamily of domains that bind organic phosphates. BMC Genomics 3:33.

30. Martinez J, Truffault V, Hothorn M. 2015. Structural Determinants for Substrate Binding and Catalysis in Triphosphate Tunnel Metalloenzymes. J Biol Chem 290:23348–60.

31. Sismeiro O, Trotot P, Biville F, Vivares C, Danchin A. 1998. Aeromonas hydrophila adenylyl cyclase 2: a new class of adenylyl cyclases with thermophilic properties and sequence similarities to proteins from hyperthermophilic archaebacteria. J Bacteriol 180:3339–44.

32. Gallagher DT, Kim SK, Robinson H, Reddy PT. 2011. Active-site structure of class IV adenylyl cyclase and transphyletic mechanism. J Mol Biol 405:787–803.

33. Gallagher DT, Smith NN, Kim SK, Heroux A, Robinson H, Reddy PT. 2006. Structure of the class IV adenylyl cyclase reveals a novel fold. J Mol Biol 362:114–22.

34. Smith N, Kim SK, Reddy PT, Gallagher DT. 2006. Crystallization of the class IV adenylyl cyclase from Yersinia pestis. Acta Crystallogr Sect F Struct Biol Cryst Commun 62:200–4.

35. Vogt MS, Ngouoko Nguepbeu RR, Mohr MKF, Albers SV, Essen LO, Banerjee A. 2021. The archaeal triphosphate tunnel metalloenzyme SaTTM defines structural determinants for the diverse activities in the CYTH protein family. J Biol Chem 297:100820.

36. Kovacikova G, Skorupski K. 2001. Overlapping binding sites for the virulence gene regulators AphA, AphB and cAMP-CRP at the *Vibrio cholerae* tcpPH promoter. Mol Microbiol 41:393–407.

37. Li CC, Merrell DS, Camilli A, Kaper JB. 2002. ToxR interferes with CRP- dependent transcriptional activation of ompT in *Vibrio cholerae*. Mol Microbiol 43:1577–89.

38. Liang W, Pascual-Montano A, Silva AJ, Benitez JA. 2007. The cyclic AMP receptor protein modulates quorum sensing, motility and multiple genes that affect intestinal colonization in *Vibrio cholerae*. Microbiology (Reading) 153:2964–2975.

39. Popovych N, Tzeng SR, Tonelli M, Ebright RH, Kalodimos CG. 2009. Structural basis for cAMP-mediated allosteric control of the catabolite activator protein. Proc Natl Acad Sci U S A 106:6927–32.

40. Shimada T, Fujita N, Yamamoto K, Ishihama A. 2011. Novel roles of cAMP receptor protein (CRP) in regulation of transport and metabolism of carbon sources. PLoS One 6:e20081.

41. Skorupski K, Taylor RK. 1997. Cyclic AMP and its receptor protein negatively regulate the coordinate expression of cholera toxin and toxin-coregulated pilus in *Vibrio cholerae*. Proc Natl Acad Sci U S A 94:265–70.

42. Whitaker WB, Richards GP, Boyd EF. 2014. Loss of sigma factor RpoN increases intestinal colonization of *Vibrio parahaemolyticus* in an adult mouse model. Infect Immun 82:544–56.

43. Kalburge SS, Carpenter MR, Rozovsky S, Boyd EF. 2017. Quorum Sensing Regulators Are Required for Metabolic Fitness in *Vibrio parahaemolyticus*. Infect Immun 85.

44. Tague JG, Hong J, Kalburge SS, Boyd EF. 2022. Regulatory Small RNA Qrr2 Is Expressed Independently of Sigma Factor-54 and Can Function as the Sole Qrr Small RNA To Control Quorum Sensing in *Vibrio parahaemolyticus*. J Bacteriol 204:e0035021.

45. Tague JG, Regmi A, Gregory GJ, Boyd EF. 2021. Fis Connects Two Sensory Pathways, Quorum Sensing and Surface Sensing, to Control Motility in *Vibrio parahaemolyticus*. Front Microbiol 12:669447.

46. McCarter LL. 1998. OpaR, a homolog of Vibrio harveyi LuxR, controls opacity of *Vibrio parahaemolyticus*. J Bacteriol 180:3166–73.

47. Guvener ZT, McCarter LL. 2003. Multiple regulators control capsular polysaccharide production in *Vibrio parahaemolyticus*. J Bacteriol 185:5431–41.

48. Enos-Berlage JL, Guvener ZT, Keenan CE, McCarter LL. 2005. Genetic determinants of biofilm development of opaque and translucent *Vibrio parahaemolyticus*. Mol Microbiol 55:1160–82.

49. Kernell Burke A, Guthrie LT, Modise T, Cormier G, Jensen RV, McCarter LL, Stevens AM. 2015. OpaR controls a network of downstream transcription factors in *Vibrio parahaemolyticus* BB22OP. PLoS One 10:e0121863.

50. Gode-Potratz CJ, McCarter LL. 2011. Quorum sensing and silencing in *Vibrio parahaemolyticus*. J Bacteriol 193:4224–37.

51. Hufnagel DA, Evans ML, Greene SE, Pinkner JS, Hultgren SJ, Chapman MR. 2016. The Catabolite Repressor Protein-Cyclic AMP Complex Regulates csgD and Biofilm Formation in Uropathogenic *Escherichia coli*. J Bacteriol 198:3329–3334.

52. Kalivoda EJ, Stella NA, O’Dee DM, Nau GJ, Shanks RM. 2008. The cyclic AMP- dependent catabolite repression system of Serratia marcescens mediates biofilm formation through regulation of type 1 fimbriae. Appl Environ Microbiol 74:3461–70.

53. Kim YR, Kim SY, Kim CM, Lee SE, Rhee JH. 2005. Essential role of an adenylate cyclase in regulating *Vibrio vulnificus* virulence. FEMS Microbiol Lett 243:497–503.

54. Kim YR, Lee SE, Kim B, Choy H, Rhee JH. 2013. A dual regulatory role of cyclic adenosine monophosphate receptor protein in various virulence traits of *Vibrio vulnificus*. Microbiol Immunol 57:273–80.

55. Charania MA, Brockman KL, Zhang Y, Banerjee A, Pinchuk GE, Fredrickson JK, Beliaev AS, Saffarini DA. 2009. Involvement of a membrane-bound class III adenylate cyclase in regulation of anaerobic respiration in *Shewanella oneidensis* MR-1. J Bacteriol 191:4298–306.

56. Fong JCN, Yildiz FH. 2008. Interplay between Cyclic AMP-Cyclic AMP Receptor Protein and Cyclic di-GMP Signaling in *Vibrio cholerae* Biofilm Formation. J Bacteriol 190:6646–6659.

57. Hammer BK, Bassler BL. 2003. Quorum sensing controls biofilm formation in *Vibrio cholerae*. Mol Microbiol 50:101–4.

58. Zhu J, Mekalanos JJ. 2003. Quorum sensing-dependent biofilms enhance colonization in *Vibrio cholerae*. Dev Cell 5:647–56.

59. Yildiz FH, Liu XS, Heydorn A, Schoolnik GK. 2004. Molecular analysis of rugosity in a *Vibrio cholerae* O1 El Tor phase variant. Mol Microbiol 53:497–515.

60. Silva AJ, Pham K, Benitez JA. 2003. Haemagglutinin/protease expression and mucin gel penetration in El Tor biotype *Vibrio cholerae*. Microbiology 149:1883–1891.

61. Ho SN, Hunt HD, Horton RM, Pullen JK, Pease LR. 1989. Site-directed mutagenesis by overlap extension using the polymerase chain reaction. Gene 77:51–9.

62. Grote A, Klein J, Retter I, Haddad I, Behling S, Bunk B, Biegler I, Yarmolinetz S, Jahn D, Münch R. 2009. PRODORIC (release 2009): a database and tool platform for the analysis of gene regulation in prokaryotes. Nucleic Acids Res 37:D61–5.

63. Crooks GE, Hon G, Chandonia JM, Brenner SE. 2004. WebLogo: a sequence logo generator. Genome Res 14:1188–90.

64. Baba T, Ara T, Hasegawa M, Takai Y, Okumura Y, Baba M, Datsenko KA, Tomita M, Wanner BL, Mori H. 2006. Construction of *Escherichia coli* K-12 in-frame, single-gene knockout mutants: the Keio collection. Mol Syst Biol 2:2006.0008.

65. Thompson JD, Higgins DG, Gibson TJ. 1994. CLUSTAL W: improving the sensitivity of progressive multiple sequence alignment through sequence weighting, position-specific gap penalties and weight matrix choice. Nucleic Acids Res 22:4673–80.

66. Kimura M. 1980. A simple method for estimating evolutionary rates of base substitutions through comparative studies of nucleotide sequences. J Mol Evol 16:111–20.

67. Kumar S, Stecher G, Li M, Knyaz C, Tamura K. 2018. MEGA X: Molecular Evolutionary Genetics Analysis across Computing Platforms. Mol Biol Evol 35:1547–1549.

68. Makino K, Oshima K, Kurokawa K, Yokoyama K, Uda T, Tagomori K, Iijima Y, Najima M, Nakano M, Yamashita A, Kubota Y, Kimura S, Yasunaga T, Honda T, Shinagawa H, Hattori M, Iida T. 2003. Genome sequence of *Vibrio parahaemolyticus*: a pathogenic mechanism distinct from that of *V. cholerae*. Lancet 361:743–9.

69. Baba T, Ara T, Hasegawa M, Takai Y, Okumura Y, Baba M, Datsenko KA, Tomita M, Wanner BL, Mori H. 2006. Construction of *Escherichia coli* K-12 in-frame, single-gene knockout mutants: the Keio collection. Mol Syst Biol 2:2006 0008.

70. Dehio C, Meyer M. 1997. Maintenance of broad-host-range incompatibility group P and group Q plasmids and transposition of Tn5 in Bartonella henselae following conjugal plasmid transfer from *Escherichia coli*. J Bacteriol 179:538–40.

71. Philippe N, Alcaraz J, Coursange E, Geiselmann J, Schneider D. 1994. Improvement of pCVD442, a suicide plasmid for gene allele exchange in bacteria. Plasmid 51:246–255.

72. Kovach ME, Phillips RW, Elzer PH, Roop RM, Peterson KM. 1994. pBBR1MCS: a broad-host-range cloning vector. Biotechniques 16:800–802.

73. Guzman LM, Belin D, Carson MJ, Beckwith J. 1995. Tight regulation, modulation, and high-level expression by vectors containing the arabinose PBAD promoter. J Bacteriol 177:4121–30.

