## Supplemental Figures for "A class IV adenylate cyclase CyaB is required for capsule polysaccharide production and biofilm formation in *Vibrio parahaemolyticus*"

### Slide 1
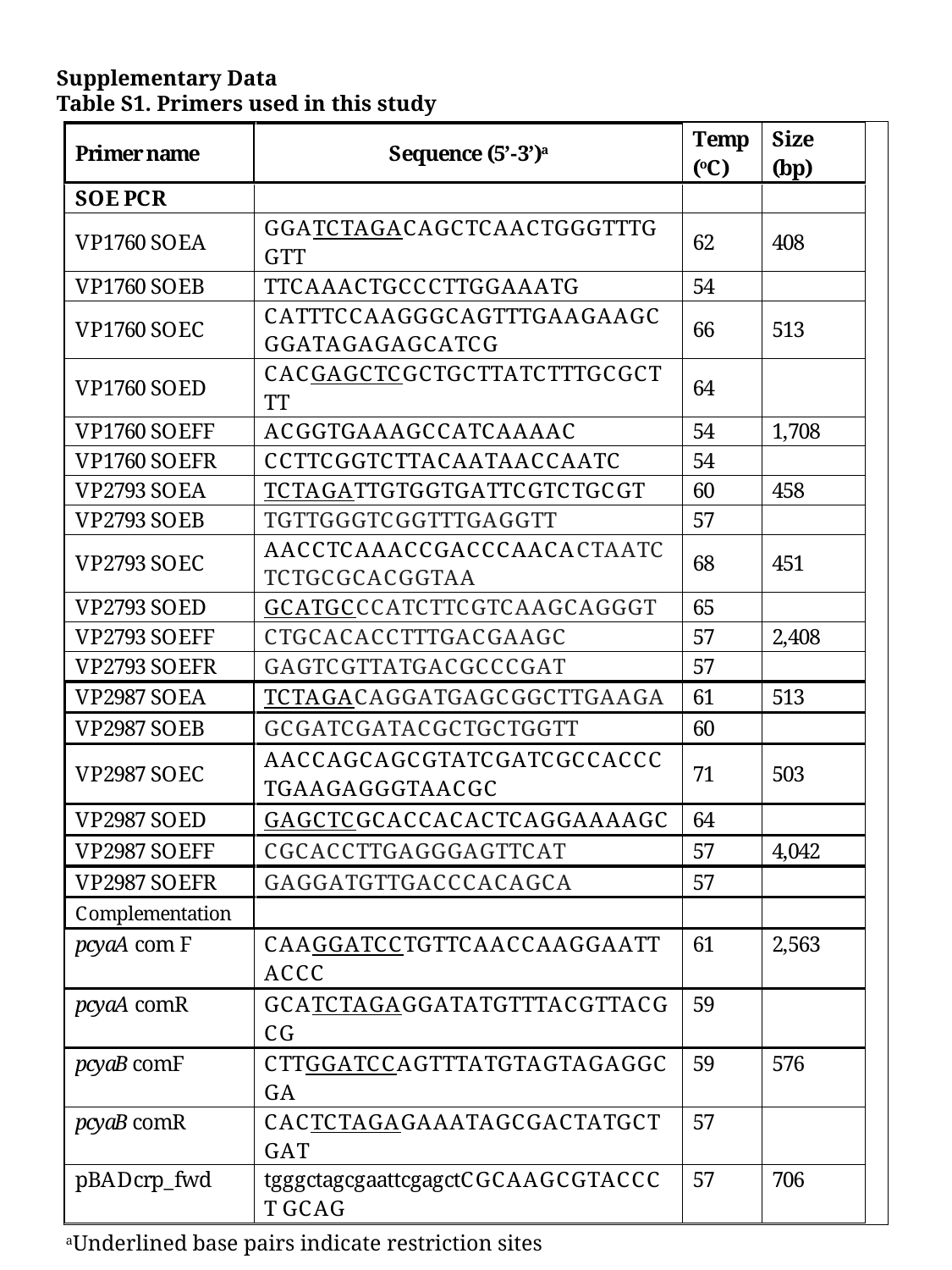

Supplementary Data
Table S1. Primers used in this study
^a^Underlined base pairs indicate restriction sites

### Slide 2
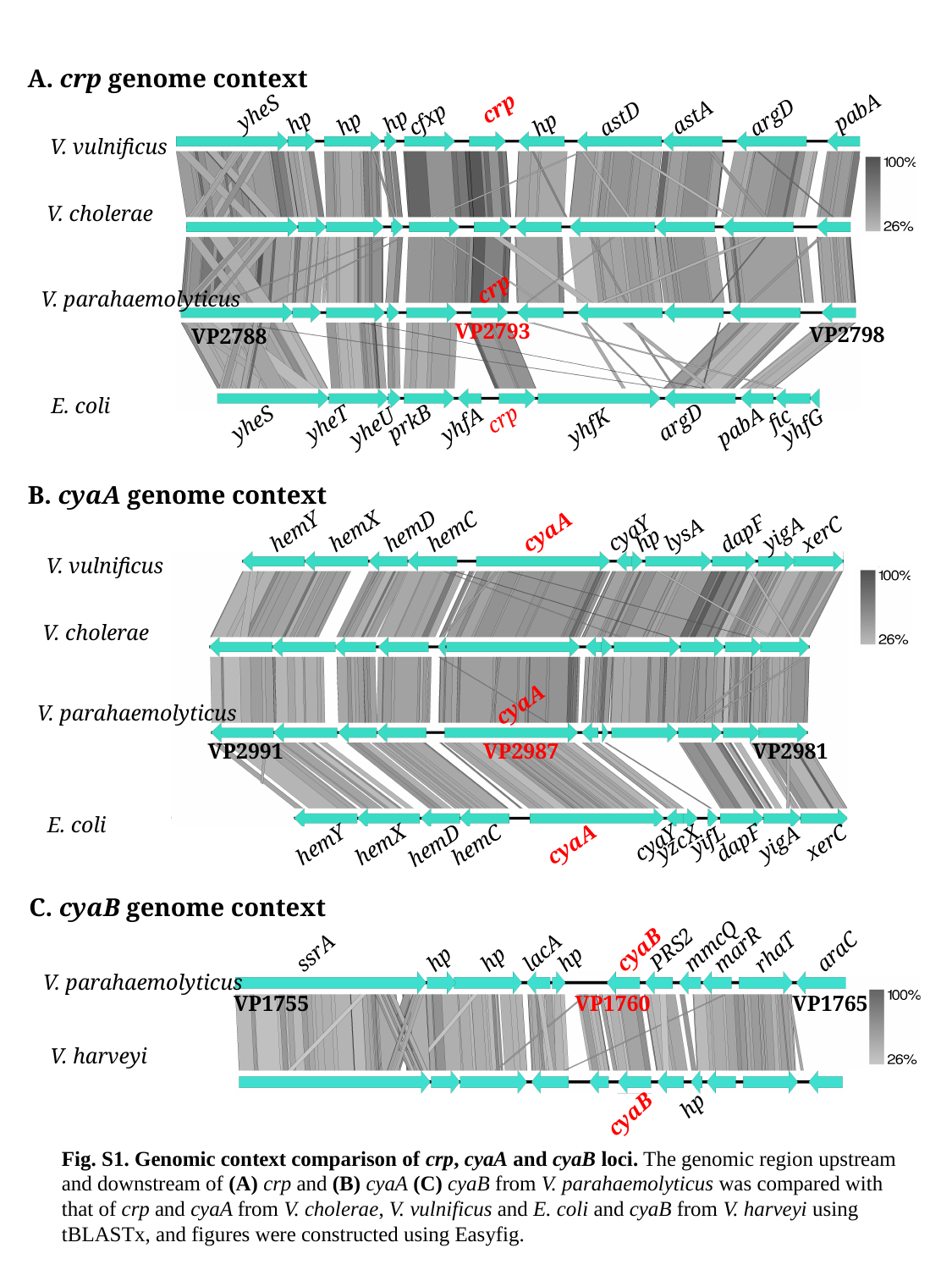

A. crp genome context
crp
pabA
yheS
argD
astA
astD
cfxp
hp
hp
hp
hp
V. vulnificus
V. cholerae
crp
V. parahaemolyticus
VP2793
VP2798
VP2788
E. coli
crp
fic
argD
prkB
yheS
yheT
yhfA
yhfK
yhfG
pabA
yheU
B. cyaA genome context
hemD
hemX
hemY
hemC
cyaA
cyaY
dapF
yigA
xerC
lysA
hp
V. vulnificus
V. cholerae
cyaA
V. parahaemolyticus
VP2991
VP2987
VP2981
E. coli
yifL
xerC
dapF
yigA
cyaY
yzcX
cyaA
hemY
hemC
hemX
hemD
C. cyaB genome context
mmcQ
PRS2
marR
cyaB
araC
rhaT
ssrA
lacA
hp
hp
hp
V. parahaemolyticus
VP1755
VP1760
VP1765
V. harveyi
hp
cyaB
Fig. S1. Genomic context comparison of crp, cyaA and cyaB loci. The genomic region upstream and downstream of (A) crp and (B) cyaA (C) cyaB from V. parahaemolyticus was compared with that of crp and cyaA from V. cholerae, V. vulnificus and E. coli and cyaB from V. harveyi using tBLASTx, and figures were constructed using Easyfig.

### Slide 3
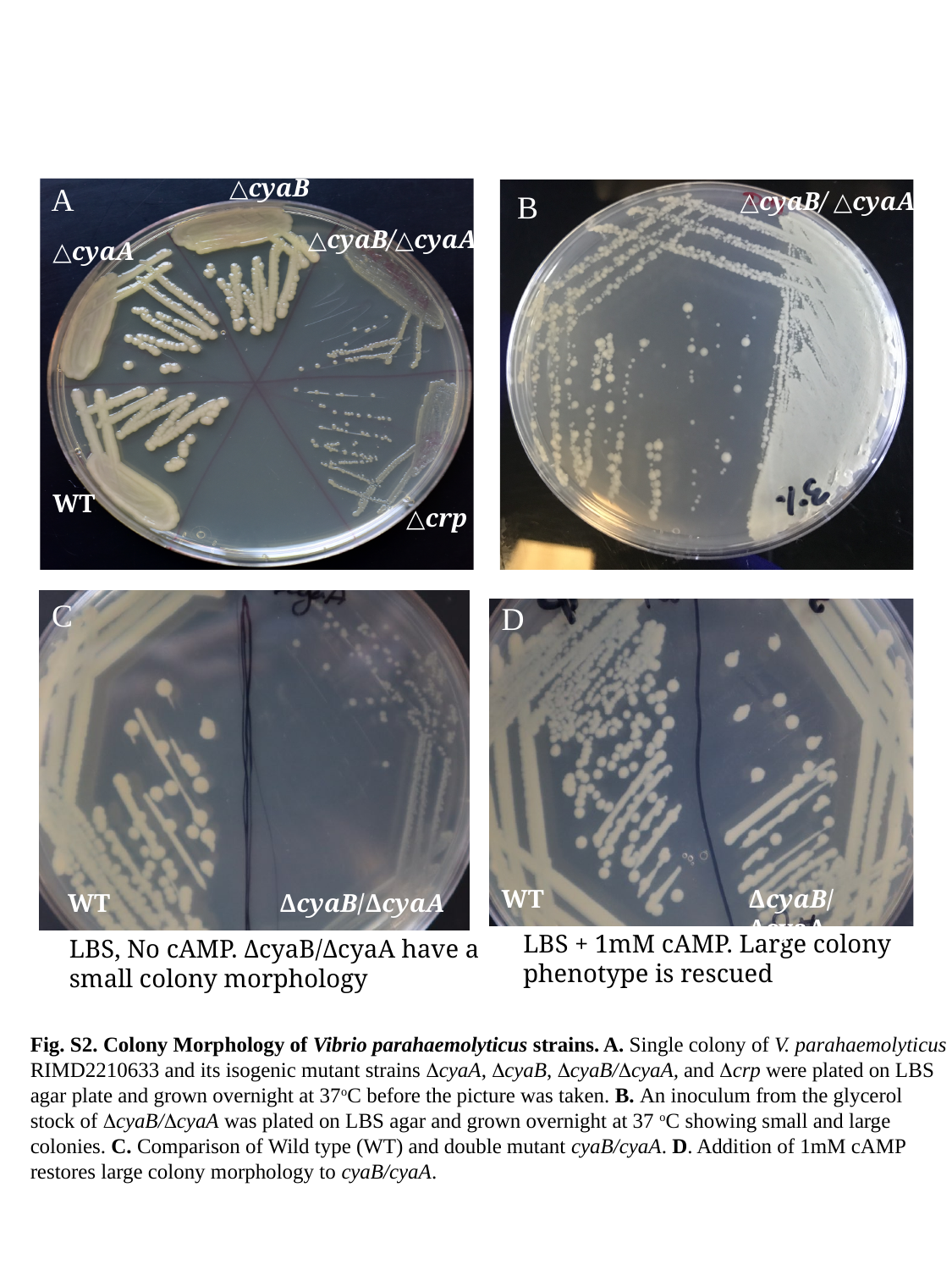

△cyaB
△cyaB/△cyaA
△cyaA
△crp
RIMD
WT
uninnoculated
△crp
A
△cyaB/ △cyaA
B
C
D
WT
ΔcyaB/ΔcyaA
WT
ΔcyaB/ΔcyaA
LBS + 1mM cAMP. Large colony phenotype is rescued
LBS, No cAMP. ΔcyaB/ΔcyaA have a small colony morphology
Fig. S2. Colony Morphology of Vibrio parahaemolyticus strains. A. Single colony of V. parahaemolyticus RIMD2210633 and its isogenic mutant strains ΔcyaA, ΔcyaB, ΔcyaB/ΔcyaA, and Δcrp were plated on LBS agar plate and grown overnight at 37^o^C before the picture was taken. B. An inoculum from the glycerol stock of ΔcyaB/ΔcyaA was plated on LBS agar and grown overnight at 37 ^o^C showing small and large colonies. C. Comparison of Wild type (WT) and double mutant cyaB/cyaA. D. Addition of 1mM cAMP restores large colony morphology to cyaB/cyaA.

### Slide 4
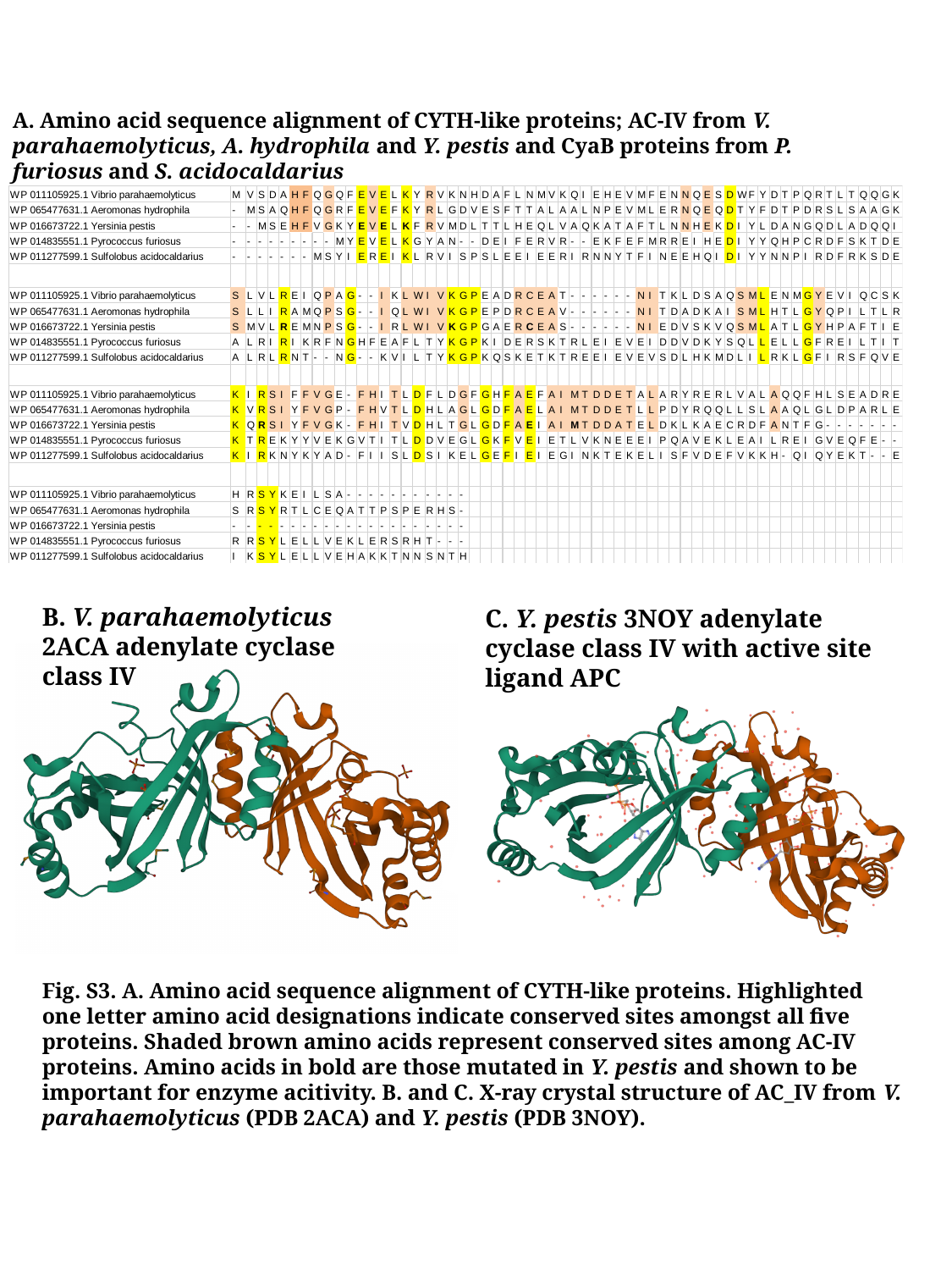

A. Amino acid sequence alignment of CYTH-like proteins; AC-IV from V. parahaemolyticus, A. hydrophila and Y. pestis and CyaB proteins from P. furiosus and S. acidocaldarius
B. V. parahaemolyticus 2ACA adenylate cyclase class IV
C. Y. pestis 3NOY adenylate cyclase class IV with active site ligand APC
Fig. S3. A. Amino acid sequence alignment of CYTH-like proteins. Highlighted one letter amino acid designations indicate conserved sites amongst all five proteins. Shaded brown amino acids represent conserved sites among AC-IV proteins. Amino acids in bold are those mutated in Y. pestis and shown to be important for enzyme acitivity. B. and C. X-ray crystal structure of AC_IV from V. parahaemolyticus (PDB 2ACA) and Y. pestis (PDB 3NOY).

### Slide 5
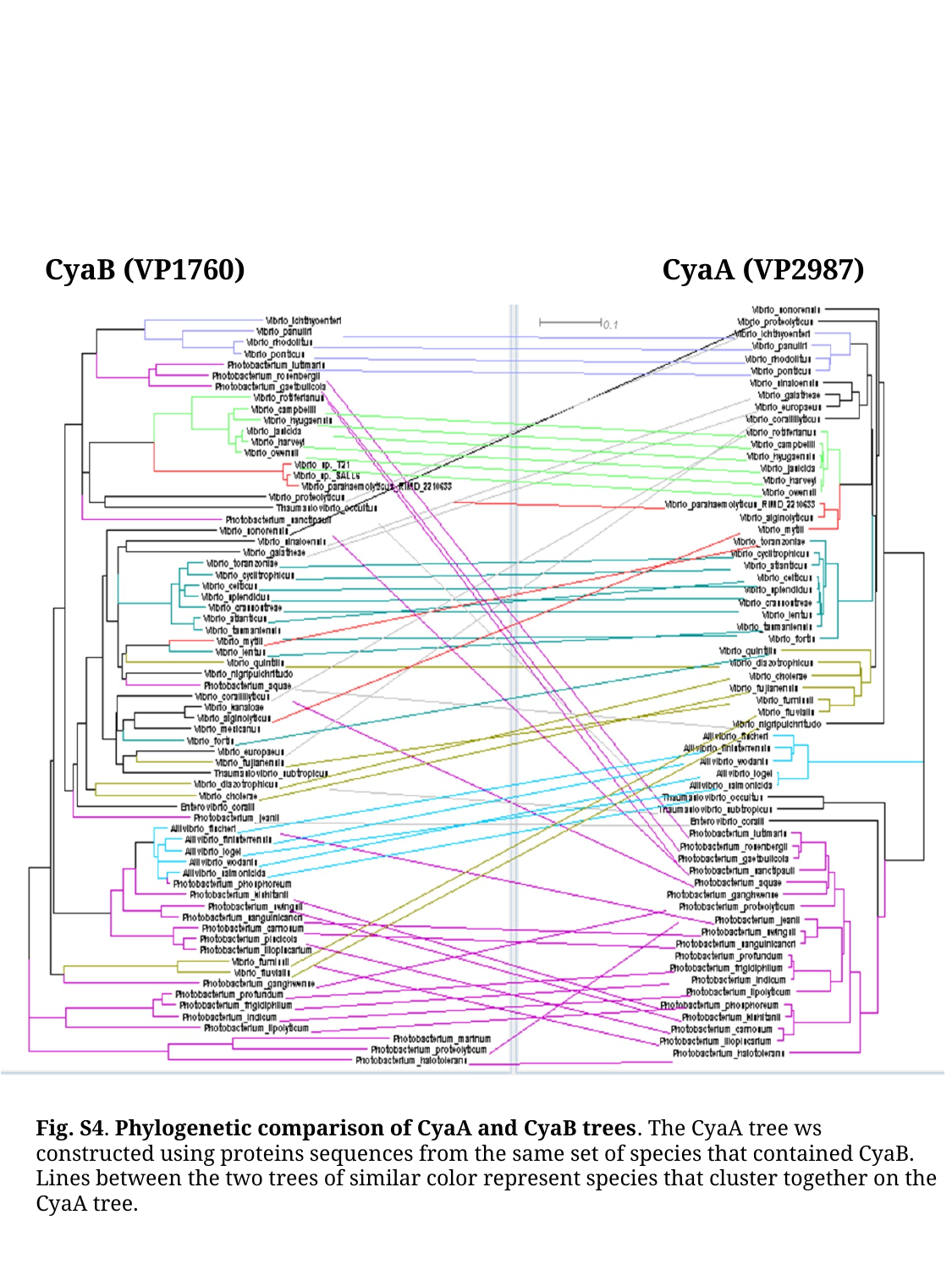

CyaB (VP1760)
CyaA (VP2987)
Fig. S4. Phylogenetic comparison of CyaA and CyaB trees. The CyaA tree ws constructed using proteins sequences from the same set of species that contained CyaB. Lines between the two trees of similar color represent species that cluster together on the CyaA tree.

### Slide 6
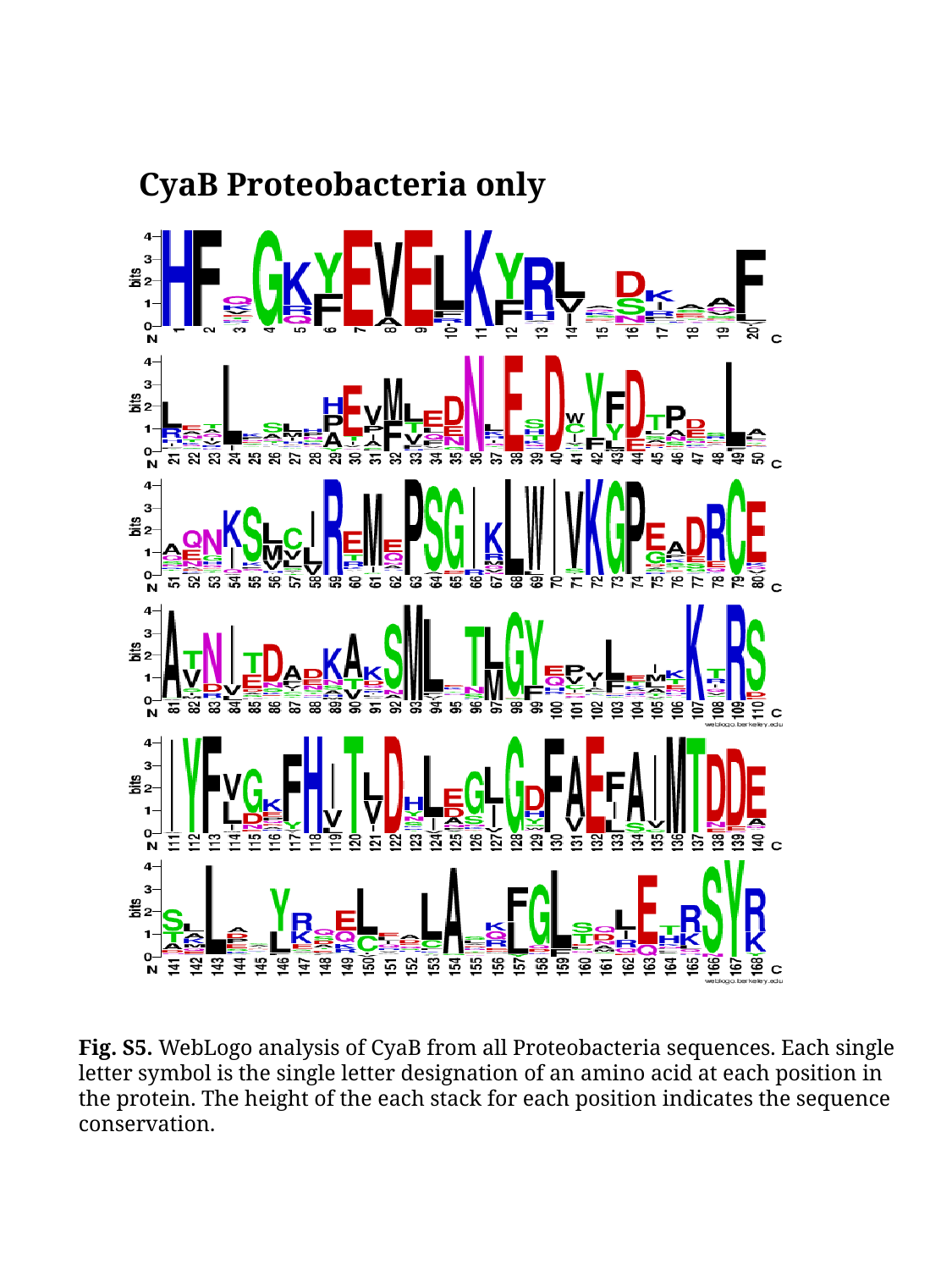

CyaB Proteobacteria only
Fig. S5. WebLogo analysis of CyaB from all Proteobacteria sequences. Each single letter symbol is the single letter designation of an amino acid at each position in the protein. The height of the each stack for each position indicates the sequence conservation.

### Slide 7
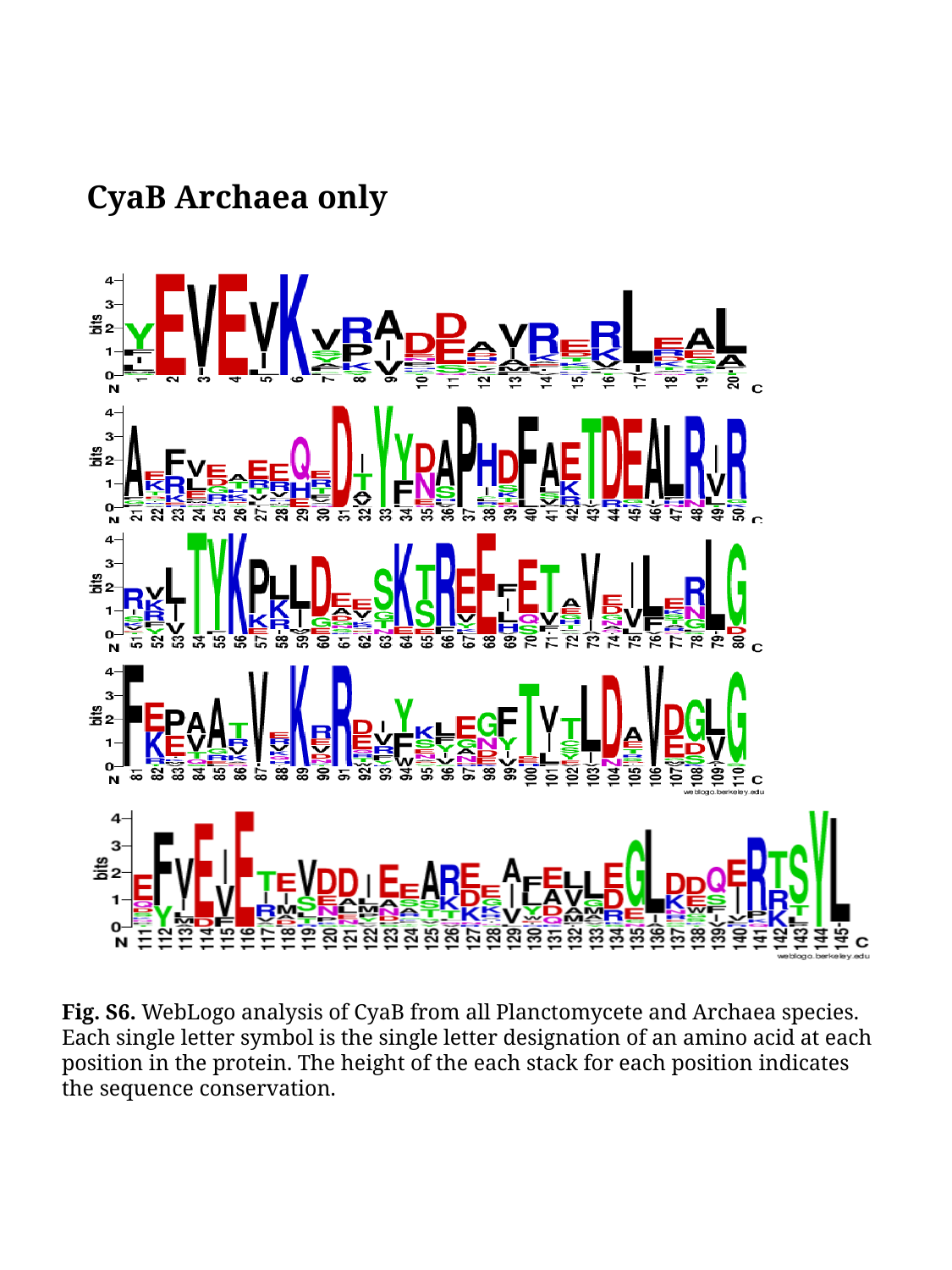

CyaB Archaea only
Fig. S6. WebLogo analysis of CyaB from all Planctomycete and Archaea species. Each single letter symbol is the single letter designation of an amino acid at each position in the protein. The height of the each stack for each position indicates the sequence conservation.

### Slide 8
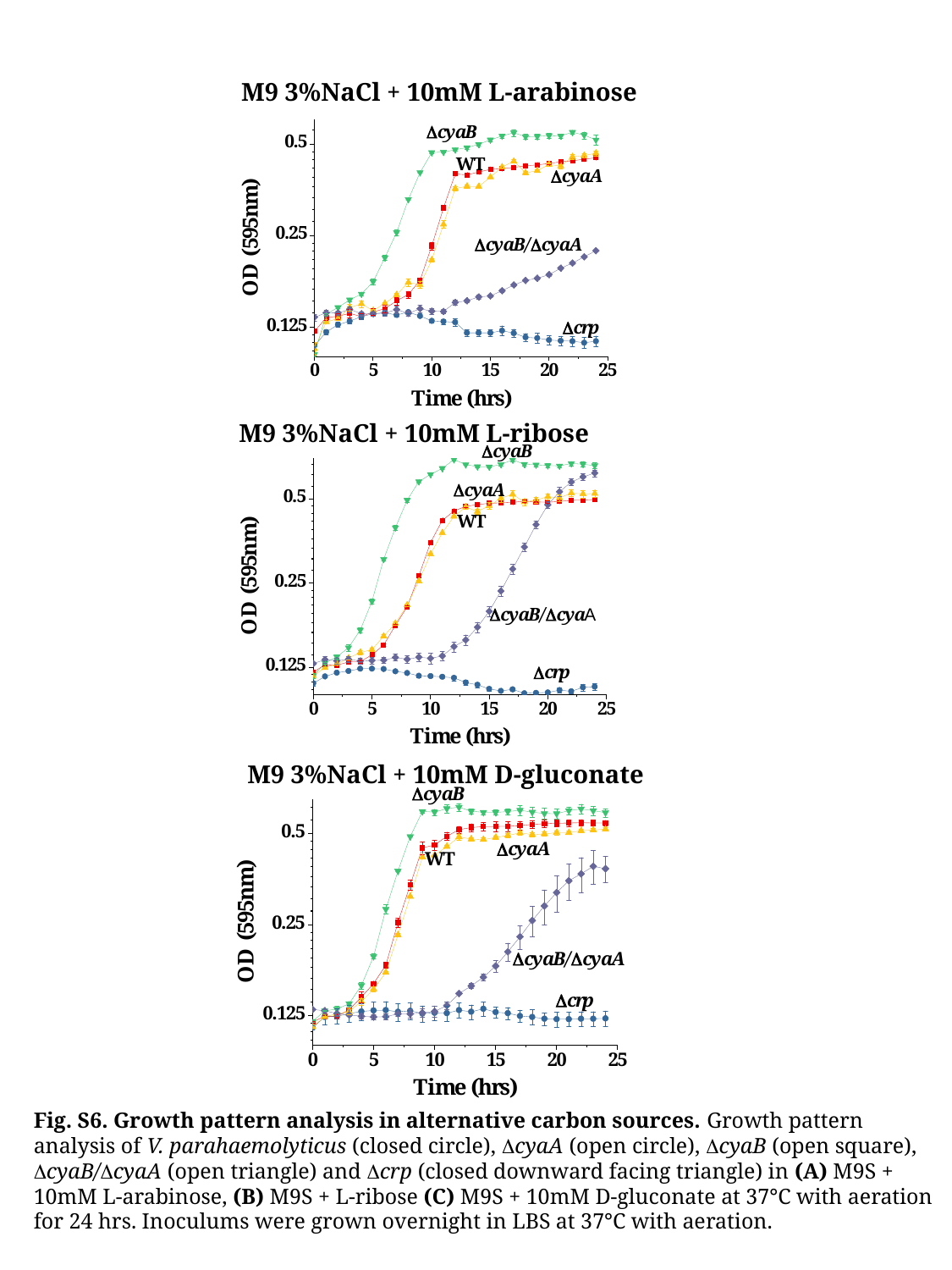

M9 3%NaCl + 10mM L-arabinose
M9 3%NaCl + 10mM L-ribose
M9 3%NaCl + 10mM D-gluconate
Fig. S6. Growth pattern analysis in alternative carbon sources. Growth pattern analysis of V. parahaemolyticus (closed circle), ΔcyaA (open circle), ΔcyaB (open square), ΔcyaB/ΔcyaA (open triangle) and Δcrp (closed downward facing triangle) in (A) M9S + 10mM L-arabinose, (B) M9S + L-ribose (C) M9S + 10mM D-gluconate at 37°C with aeration for 24 hrs. Inoculums were grown overnight in LBS at 37°C with aeration.

### Slide 9
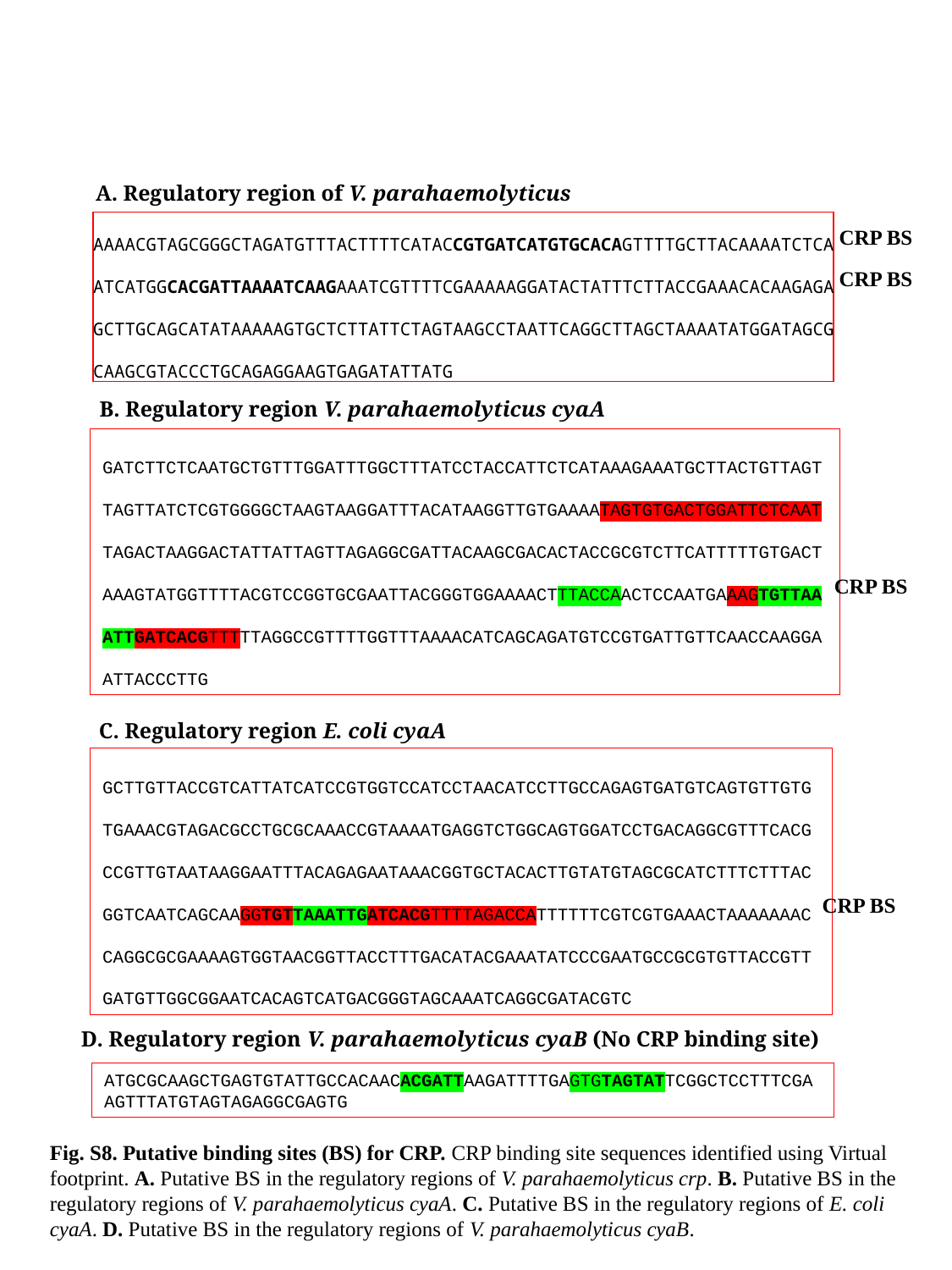

A. Regulatory region of V. parahaemolyticus crp
AAAACGTAGCGGGCTAGATGTTTACTTTTCATACCGTGATCATGTGCACAGTTTTGCTTACAAAATCTCA ATCATGGCACGATTAAAATCAAGAAATCGTTTTCGAAAAAGGATACTATTTCTTACCGAAACACAAGAGA GCTTGCAGCATATAAAAAGTGCTCTTATTCTAGTAAGCCTAATTCAGGCTTAGCTAAAATATGGATAGCG CAAGCGTACCCTGCAGAGGAAGTGAGATATTATG
CRP BS
CRP BS
B. Regulatory region V. parahaemolyticus cyaA
GATCTTCTCAATGCTGTTTGGATTTGGCTTTATCCTACCATTCTCATAAAGAAATGCTTACTGTTAGTTAGTTATCTCGTGGGGCTAAGTAAGGATTTACATAAGGTTGTGAAAATAGTGTGACTGGATTCTCAATTAGACTAAGGACTATTATTAGTTAGAGGCGATTACAAGCGACACTACCGCGTCTTCATTTTTGTGACTAAAGTATGGTTTTACGTCCGGTGCGAATTACGGGTGGAAAACTTTACCAACTCCAATGAAAGTGTTAAATTGATCACGTTTTTAGGCCGTTTTGGTTTAAAACATCAGCAGATGTCCGTGATTGTTCAACCAAGGAATTACCCTTG
CRP BS
C. Regulatory region E. coli cyaA
GCTTGTTACCGTCATTATCATCCGTGGTCCATCCTAACATCCTTGCCAGAGTGATGTCAGTGTTGTGTGAAACGTAGACGCCTGCGCAAACCGTAAAATGAGGTCTGGCAGTGGATCCTGACAGGCGTTTCACGCCGTTGTAATAAGGAATTTACAGAGAATAAACGGTGCTACACTTGTATGTAGCGCATCTTTCTTTAC
GGTCAATCAGCAAGGTGTTAAATTGATCACGTTTTAGACCATTTTTTCGTCGTGAAACTAAAAAAAC
CAGGCGCGAAAAGTGGTAACGGTTACCTTTGACATACGAAATATCCCGAATGCCGCGTGTTACCGTTGATGTTGGCGGAATCACAGTCATGACGGGTAGCAAATCAGGCGATACGTC
CRP BS
D. Regulatory region V. parahaemolyticus cyaB (No CRP binding site)
ATGCGCAAGCTGAGTGTATTGCCACAACACGATTAAGATTTTGAGTGTAGTATTCGGCTCCTTTCGAAGTTTATGTAGTAGAGGCGAGTG
Fig. S8. Putative binding sites (BS) for CRP. CRP binding site sequences identified using Virtual footprint. A. Putative BS in the regulatory regions of V. parahaemolyticus crp. B. Putative BS in the regulatory regions of V. parahaemolyticus cyaA. C. Putative BS in the regulatory regions of E. coli cyaA. D. Putative BS in the regulatory regions of V. parahaemolyticus cyaB.

### Slide 10
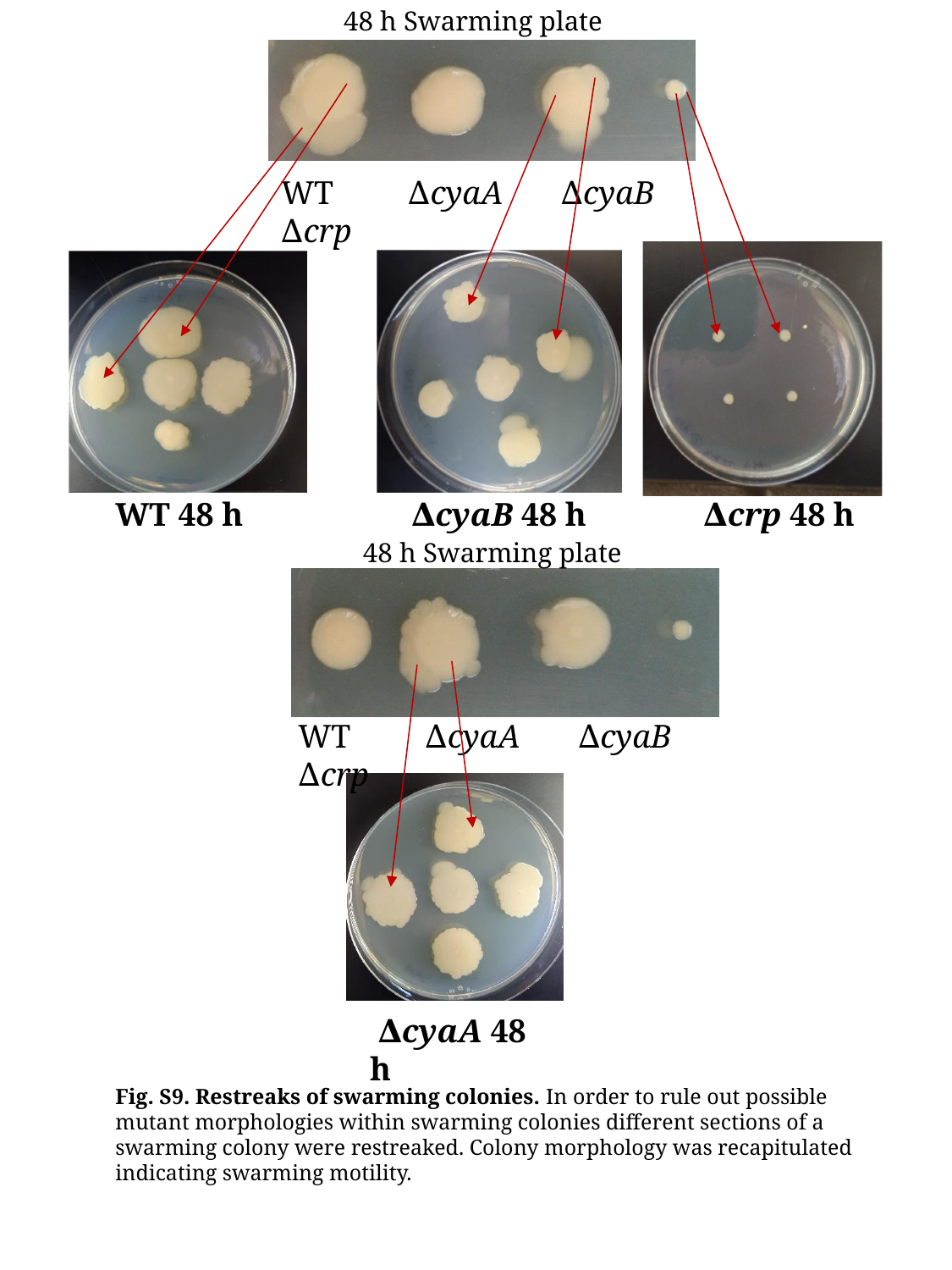

48 h Swarming plate
WT ∆cyaA ∆cyaB ∆crp
WT 48 h
 ∆cyaB 48 h
 ∆crp 48 h
48 h Swarming plate
WT ∆cyaA ∆cyaB ∆crp
 ∆cyaA 48 h
Fig. S9. Restreaks of swarming colonies. In order to rule out possible mutant morphologies within swarming colonies different sections of a swarming colony were restreaked. Colony morphology was recapitulated indicating swarming motility.

### Slide 11
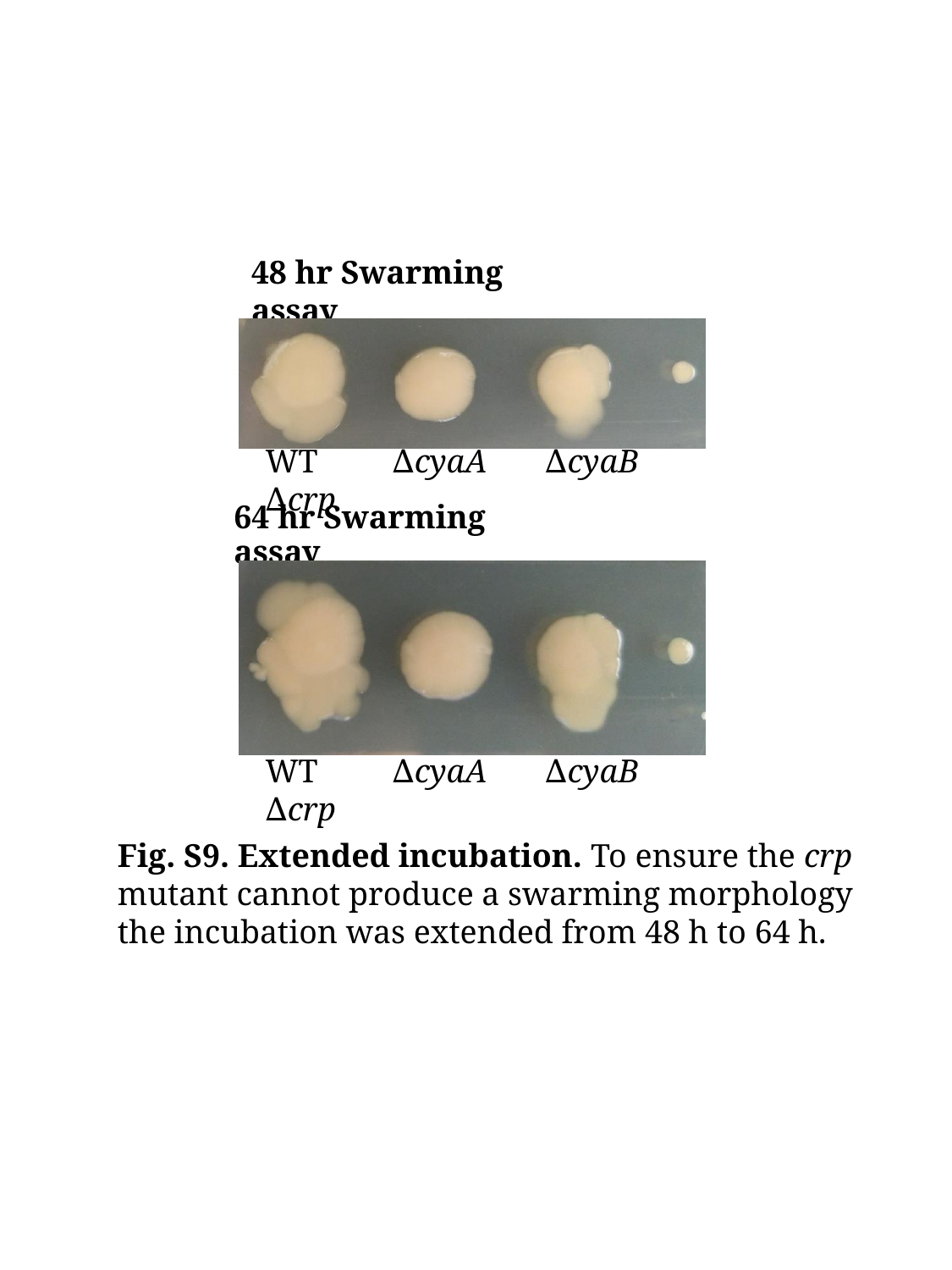

#### 48 hr Swarming assay
WT ∆cyaA ∆cyaB ∆crp
64 hr Swarming assay
WT ∆cyaA ∆cyaB ∆crp
Fig. S9. Extended incubation. To ensure the crp mutant cannot produce a swarming morphology the incubation was extended from 48 h to 64 h.

### Slide 12
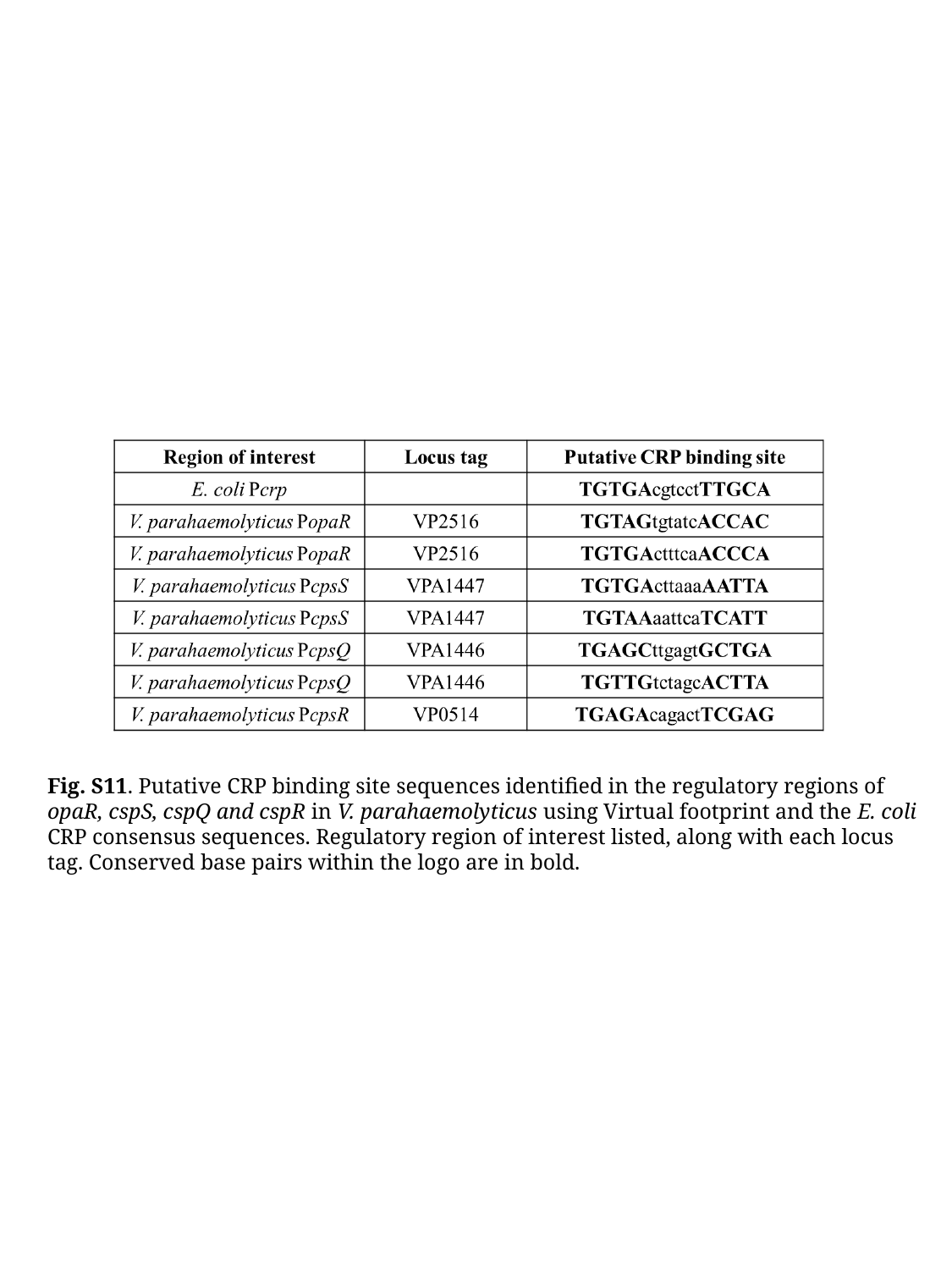

Fig. S11. Putative CRP binding site sequences identified in the regulatory regions of opaR, cspS, cspQ and cspR in V. parahaemolyticus using Virtual footprint and the E. coli CRP consensus sequences. Regulatory region of interest listed, along with each locus tag. Conserved base pairs within the logo are in bold.
